# Unique transcriptional profiles of adult human immature neurons in healthy aging, Alzheimer’s disease, and cognitive resilience

**DOI:** 10.1101/2025.01.08.631686

**Authors:** G Tosoni, D Ayyildiz, A Penning, S Snoeck, E Santiago-Mujika, O Ruiz Ormaechea, H Lee, SM Poovathingal, K Davie, J Bryois, W Macnair, J Anink, LE De Vries, J Verhaagen, E Aronica, S Thuret, O Basak, L Roybon, CP Fitzsimons, PJ Lucassen, E Salta

## Abstract

The existence and functional significance of immature neurons in the adult human brain, particularly in the context of neurodegenerative disorders, remain controversial. While rodent studies have highlighted active roles for adult-born immature neurons in the hippocampus under both healthy conditions and in Alzheimer’s disease (AD), evidence from the human brain is limited and lacks detailed molecular characterization. To address this gap, we performed single-nucleus RNA sequencing in aged healthy, AD and dementia-resilient human hippocampus to probe immature neuronal signatures and gene expression alterations associated with AD pathology and resilience. Employing a novel experimental and computational pipeline, we identified persistent populations of immature neurons across all donor groups, with transcriptional profiles distinct from both fetal counterparts and adult mature hippocampal neurons. These profiles were associated with ‘juvenile’ cellular functions, suggesting that the presence of these immature neuronal populations *per se* may actively contribute to maintaining homeostasis within the aged human hippocampus, a role that may be disrupted in AD. In the resilient brain, immature neurons were involved in transcriptional programs and intercellular interactions associated with anti-inflammatory, neurotrophic, neuroprotective, myelinating, anti-apoptotic and anti-amyloidogenic signaling pathways, suggesting active roles for the immature cells in enhancing cognitive resilience in the presence of AD pathology. Our findings reveal novel, putative physiological roles for immature neurons in the healthy and resilient adult human brain, and offer a resource for probing new strategies with potential functional relevance in AD.

## Introduction

An attractive approach to treating Alzheimer’s disease (AD) could involve adding endogenous ‘functional units’ to the degenerating hippocampal network^1^. This strategy presumes the occurrence of adult hippocampal neurogenesis (AHN) in the human brain and the functional integration of newly generated granule cell (GC) neurons into the hippocampal formation. Reconstructing the molecular and cellular signatures of immature hippocampal granule cells may not only offer novel targets for brain repair and regeneration strategies in the AD human brain; it can also directly probe the question of whether human AHN contributes to a lifelong buildup of cognitive reserve, which may later on confer resilience to AD-related dementia^2,3^.

Thus far, identifying and profiling putatively neurogenic populations in the adult human brain has not been trivial, which apart from reflecting a series of technical roadblocks, also hints at some potentially unique attributes of these cells^1,4–6^. The expected scarcity of neurogenic events in the adult —and particularly in the aging— human hippocampus, along with the possibly human-specific and also mixed-cell type gene expression profiles of immature neurons and progenitors that transit along a differentiation continuum, necessitate novel experimental and computational pipelines, and stringent quality control^5^.

Going beyond the rather limiting approach of using only a small set of markers to survey human brain tissue for immature neurons or progenitors, single-cell or single-nucleus RNA sequencing (sc/snRNAseq)-based studies have over the past years identified cells with immature neuronal characteristics in the adult human hippocampus^7–9^. Recently, we critically and comprehensively analyzed experimental and computational variables that may confound the results and conclusions of such approaches^4^. Constructing a consensus of putatively neurogenic signatures in the adult human brain, let alone in AD, would require overcoming data quality discrepancies and batch effects across technological platforms and studies. To achieve this, novel experimental and analytical strategies are essential^8,10,11^.

Taking these valuable lessons into consideration, we here established a new experimental and computational pipeline to, first, enrich for and reliably identify putatively neurogenic populations in the aged human brain without introducing the bias of marker-based preselection, and second, address their relevance for AD pathology and resilience. We find that immature neuronal signatures persist into adulthood. While they present some transcriptional similarities to their fetal counterparts, they may have additionally acquired unique features, likely allowing them to adapt to the complex adult niche microenvironment. Transcriptional profiling suggests that adult immature neurons may contribute to the homeostasis of the hippocampal niche in both healthy and resilient aged human brain, by exerting unique, non-canonical functions, which may be compromised in AD.

## Results

### Enrichment and transcriptomic characterization of the neurogenic niche in adult human hippocampus

We first performed single-nucleus RNA sequencing (snRNA-seq) on high quality and short postmortem delay (PMD) fresh frozen hippocampal tissue from 24 individuals (Figure 1A). Donors were grouped based on AD pathology and cognitive status: severe AD (SAD, n=7) and moderate AD (MAD, n=5), both consisting of demented individuals with confirmed AD histopathology; a control group (CTRL, n=6) of cognitively intact individuals without AD histopathology; and a resilient group (RES, n=6) of individuals who presented with AD histopathology when assessed postmortem, but remained cognitively intact before death (Figure 1B, Supplementary Table 1.1). To enhance our analytical power, we employed a targeted isolation strategy to primarily microdissect the granular cell layer (GCL) and subgranular zone (SGZ) of the dentate gyrus (DG), which have been shown to harbor neurogenic niche characteristics^2,7–9,12–14^, in order to enrich for ‘niche’-resident cell populations, including rare putative neurogenic cell types. To adopt an agnostic approach, we generated unsorted snRNAseq data, without prior marker-based cell selection. After quality control filtering, ambient RNA removal, and data harmonization, we obtained 110,527 high-quality nuclei (Supplementary Table 1.2, Supplementary Figure 1A-C). Using unsupervised clustering and well-established markers, we successfully identified all major DG cell types, including astrocytes (*ALDOC, AQP4*), oligodendrocytes (*MOBP, MAG*), oligodendrocyte progenitor cells (OPCs; *PDGFRA, OLIG1*), microglia (*CSF1R, PTPRC*), vascular leptomeningeal cells (VLMCs; *DCN, COL1A2*), endothelial cells (*CLDN5, VWF*), inhibitory neurons (InN; *GAD1, GAD2*), and excitatory neurons (ExN; *RBFOX3, SLC17A7*) (Figure 1C, D, Supplementary Table 1.3). Among the excitatory neurons, we further distinguished between mossy cells/Cornu Ammonis (MC/CA; *CFAP299, SV2B*) and granule cell (GC) neurons (*PROX1, SEMA5A*). Notably, GC, the principal neuronal population within the neurogenic niche, constituted 54% of all cells identified in our dataset, representing a substantial enrichment compared to previous snRNA-seq and scRNA-seq studies investigating AHN, where GC proportions were substantially lower: 7% in Ayhan et al.^15^, 10% in Habib et al.^16^, 19% in Yao et al.^7^, and 23% in both Franjic et al.^17^and Zhou et al.^8^ (Supplementary Figure 1D). This marked increase in GC representation underscored the effectiveness of our isolation protocol, offering enhanced power to detect transcriptomic signatures of less abundant populations compared to previous studies. The distribution of major cell types was consistent across different cognitive and pathological groups, with some apparent trends, but no significant differences (see Methods, Compositional Analysis) in relative cell type proportions (Figure 1E, Supplementary Table 1.4).

**Figure 1.**
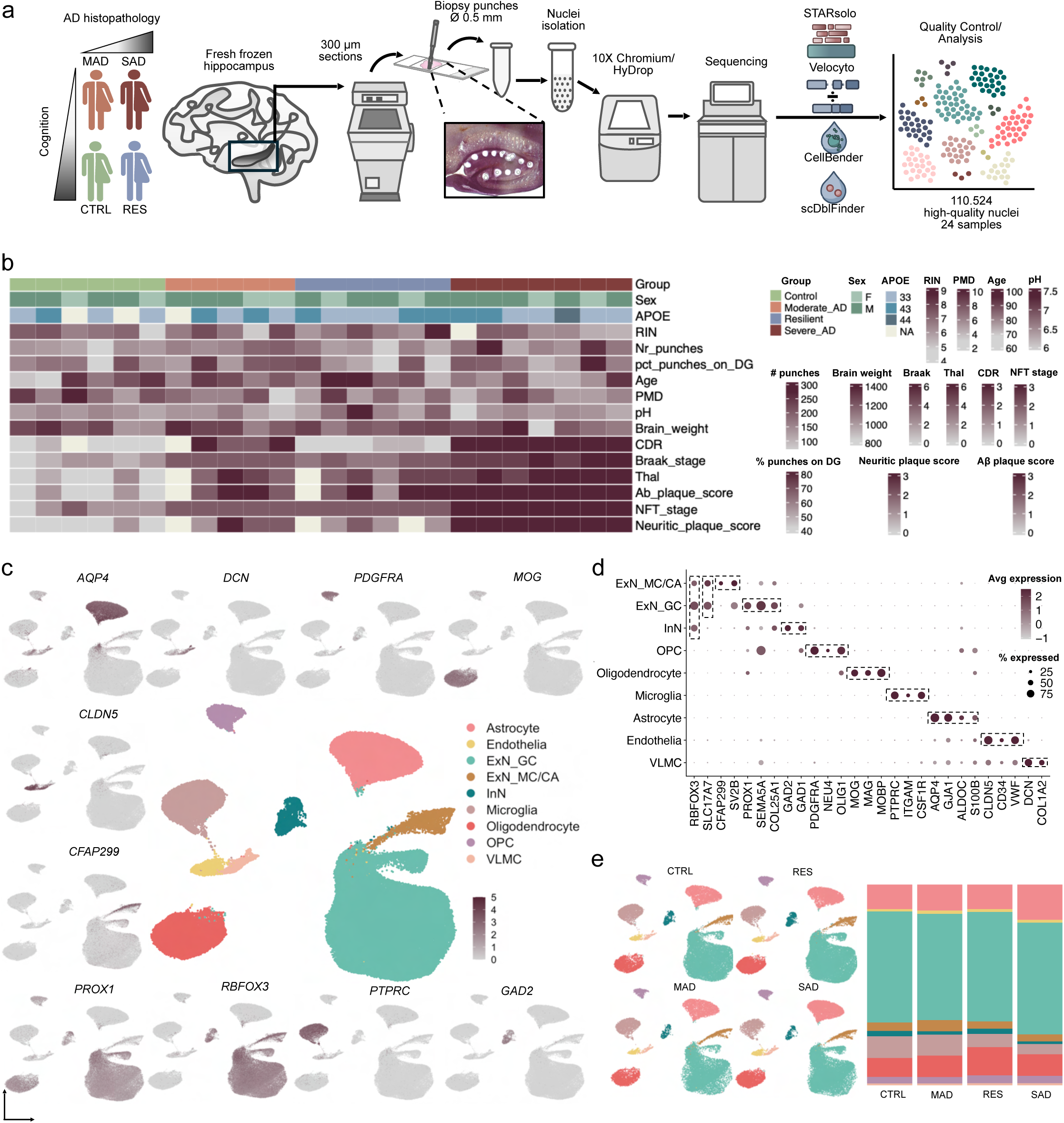
Isolation and snRNA-seq of the adult hippocampal niche. **A**, Schematic illustration of the experimental and computational design. CTRL, control; RES, resilient; MAD, moderate Alzheimer’s disease; SAD, severe Alzheimer’s disease. **B**, Cohort metadata overview. The heatmap shows sex, APOE genotype, quality (RIN) and abundance of input material (Nr punches, pct_punched on DG), age at death, postmortem delay (PMD), pH and brain weight, and measures of AD pathology and cognitive function (rows) for the 24 samples (columns), ordered by assigned group (CTRL, MAD, RES, SAD). **C**, Uniform Manifold Approximation and Projection (UMAP) plot representing 110,527 nuclei, categorized into 9 major adult hippocampal populations, and expression of representative genes for the indicated cellular populations. Astrocytes (*AQP4*, 15,277 nuclei), Endothelia (*CLDN5*, 1570 nuclei), Excitatory Neuron Granule Cells (ExN_GC, *PROX1*, *RBFOX3*, 60,461 nuclei), Excitatory Neurons Mossy Cell/Cornu Ammonis (ExN_MC/CA, *CFAP299, RBFOX3,* 4,531 nuclei), Inhibitory Neurons (InN, *GAD2,* 2,306 nuclei), Microglia (*PTPRC*, 8,723 nuclei), Oligodendrocytes (*MOG*, 12,686 nuclei), Oligodendrocyte Progenitor Cells (OPC, *PDGFRA,* 3,917 nuclei), Vascular and Leptomeningeal Cells (VLMC, *DCN,* 1,056 nuclei). **D**, Dot plot of representative genes specific for the indicated cell populations. The size of each dot represents the cell percentage of this population positive for the marker gene. The scale of the dot color represents the average expression level of the marker gene in this population. **E**, Relative proportion of each cell population in the 4 different groups (CTRL, MAD, RES, SAD) represented as UMAPs and barplot. RIN, RNA integrity number; Nr, number; pct, percentage; CDR, Clinical Dementia Rating.

### Neurogenic signatures in human fetal hippocampus

To establish a reliable ‘positive control’ reference for neurogenic profiles, we next employed fetal hippocampal tissue, where neurogenesis is abundant and easily identifiable. Developmental and adult neurogenesis were previously shown to share similar fundamental processes and transcriptomic profiles^18,19^.

We generated a dataset of 41,984 high-quality nuclei from 3 samples of human fetal hippocampal tissue at gestational week 24, a stage at which the DG is already formed^20,21^ (Figure 2A, Supplementary Table 2.1). Through unsupervised clustering and the use of canonical markers, we successfully identified all major hippocampal cell types (Figure 2B,C, Supplementary Figure 2A,B, Supplementary Table 2.2). We then focused our analysis on neurogenic populations within the DG, including astrocytes/neural stem cells (Astrocyte/NSC; *SOX9*, *HES5*), non-cycling progenitors (*HES6*, *EGFR*), cycling progenitors (*MKI67*, *TOP2A*), and neuroblasts/immature neurons (NB/ImN; *PROX1*, *EPHA7*, *SEMA3C*) (Figure 2D, Supplementary Table 2.3). Pseudotime and cell cycle analyses confirmed the presence of a neurogenic trajectory, progressing from NSC, through progenitors, to immature neurons (Figure 2E, F, Supplementary Figure 2C, Supplementary Tables 2.4, 2.5, 2.6).

**Figure 2.**
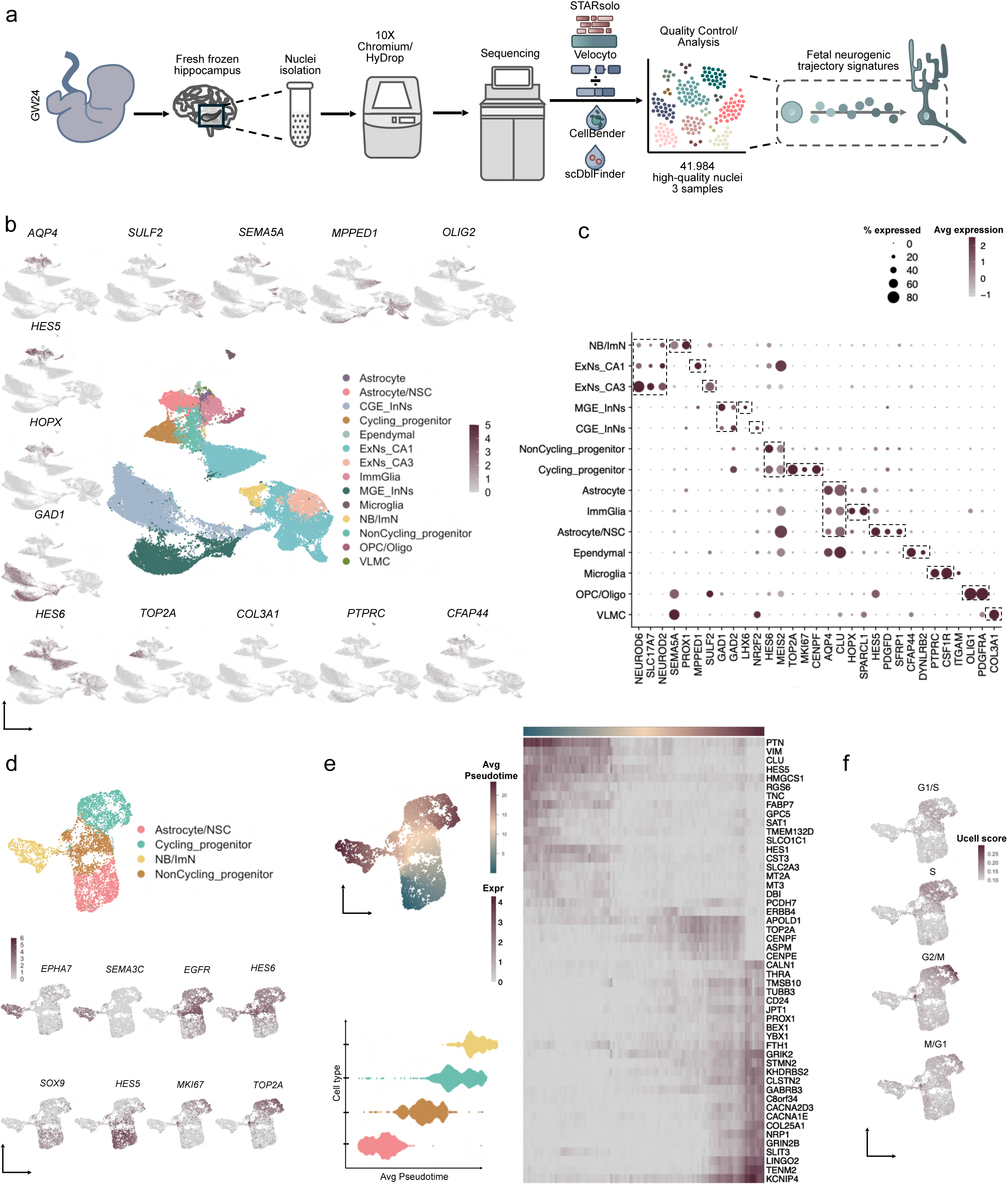
Isolation and snRNA-seq of the fetal hippocampal neurogenic niche. **A,** Schematic illustration of the experimental and computational design. **B,** UMAP plot representing 41,984 nuclei, categorized into 14 major fetal hippocampal populations, and expression of representative genes for the indicated cell populations. Astrocyte (*AQP4*, 504 nuclei), Astrocyte/Neural Stem Cell (NSC, *HES5,* 1,695 nuclei), Caudal Ganglionic Eminence Inhibitory Neurons (CGE_InN, *GAD1, NR2F2*, 14,133 nuclei), Cycling neuronal progenitor (*TOP2A*, 1,643 nuclei), Ependymal (*CFAP44*, 191 nuclei), Excitatory Neuron Cornu Ammonis 1 (ExN_CA1, *MMPED1*, 10,970 nuclei), Excitatory Neuron Cornu Ammonis 3 (ExN_CA3, *SULF2*, 1,482 nuclei), Immature Glia (ImmGlia, *HOPX*, 1,144 nuclei), Medial Ganglionic Eminence Inhibitory Neurons (MGE_InN, *GAD1, LHX6*, 7,151 nuclei), Microglia (*PTPRC*, 297 nuclei), Neuroblast/Immature Neuron (NB/ImN, *SEMA5A*, 838 nuclei), non-cycling neuronal progenitor (*HES6,* 1,438 nuclei), Oligodendrocyte Progenitor Cell/Oligodendrocyte (OPC/Oligo, *OLIG2,* 445 nuclei), Vascular and Leptomeningeal Cell (VLMC, *COL3A1,* 53 nuclei). **C,** Dot plot of representative genes for the indicated cell populations. The size of each dot represents the cell percentage of this population positive for the marker gene. The scale of the dot color represents the average expression level of the marker gene in this population. **D,** UMAP plot of the fetal neurogenic lineage and expression of representative genes specific for the indicated cell populations. **E,** Pseudotemporal ordering of cells in the neurogenic lineage visualized as UMAP and violin plot (left), and heatmap of differentially expressed genes along pseudotime (right). **F,** Cell cycle inference in the neurogenic lineage, performed using gene signature scoring, based on cell cycle phase-specific genes identified by Schwabe et al.^119^ (Supplementary Table 2.6). GW, gestational week.

### *In silico* filtering of ‘noisy’ transcriptional signatures in adult GC

Subsequently, we delved deeper into the adult GC subset, where we would expect to find putative immature neuronal signatures. Given that neurogenic cell types are part of a continuum and may thus exist in intermediate or transitioning states^22^, effectively removing ambient RNA contamination is crucial. Ambient RNA contamination poses a significant challenge in sc/snRNA-seq workflows, often distorting gene expression profiles and complicating the distinction between non-empty (cell/nuclei-containing) and empty droplets. In addition, without proper correction, ambient RNA signatures can obscure rare cell types or lead to misidentification of cell populations, falsely implying intermediate states and resulting in inaccurate interpretations^23,24^.

To address this issue, we implemented a rigorous additional filtering pipeline to the adult ExNeuron_GC population (Figure 3A, Supplementary Figure 3A-C). We first used *CellBender*^25^ to remove ambient RNA contamination. Next, to further ensure that the presence of glial markers in certain GC subtypes (Supplementary Table 3.1) reflected endogenous signatures rather than contamination artefacts, we subset adult astrocytes and GC. Re-clustering of the two populations confirmed that GC subtypes were transcriptionally distinct from astrocytes (Supplementary Figure 3A). Subsequently, we integrated the GC dataset with empty droplets that had been previously filtered out, and we removed the cells co-clustering with empty droplets (n=1386 nuclei, Supplementary Figure 3B). Then we calculated the empty-droplet RNA profile, and removed the resulting gene set (Supplementary Table 3.2) from the highly variable genes of the GC dataset. Additionally, we repeated the clustering using the unspliced RNA matrix, which reproduced the original clusters (Supplementary Figure 3C), suggesting that the observed clustering is not driven by cytoplasmic (spliced) RNA debris^24^, thereby confirming the robustness of our cell clustering and annotation strategy.

**Figure 3.**
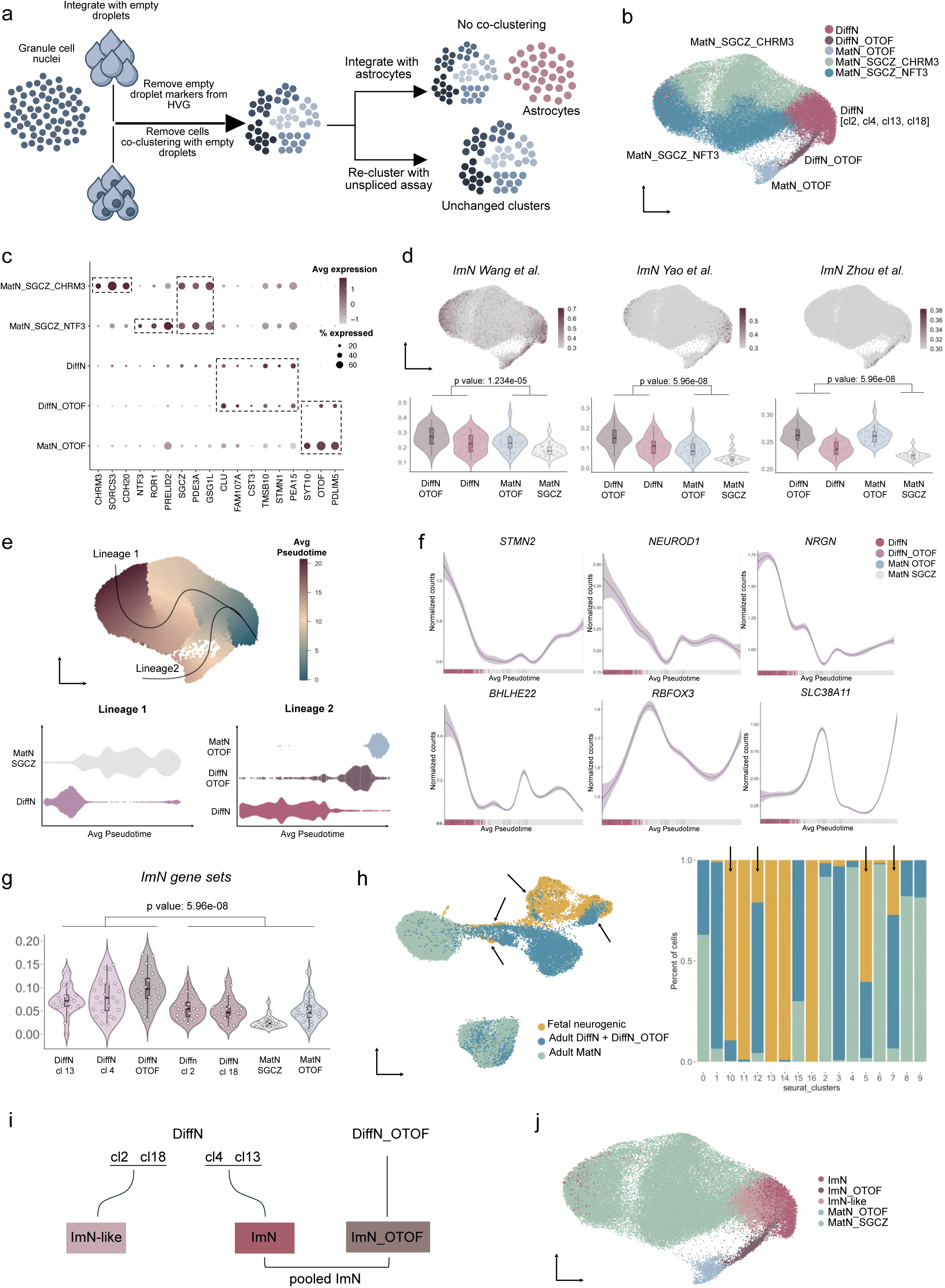
Identification of immature neurons in the adult brain. **A,** Schematic representation of the additional quality control steps performed prior to probing the GC subset for neurogenic transcriptomic signatures. **B,** UMAP plot representing 59,075 granule cell nuclei categorized into 5 subtypes. See also Supplementary Figure 3D for further information on Seurat cluster numbers. **C,** Dot plot of representative genes specific for the indicated cell populations. The size of each dot represents the cell percentage of this population positive for the marker gene. The scale of the dot color represents the average expression level of the marker gene in this population. **D**, Combined expression levels of gene groups defining the ImN transcriptional profile (as characterized in previous studies^7–9^, Supplementary Table 3.3) are represented using UMAP visualization (upper panel) and violin plots (lower panel). In the violin plots, each dot corresponds to the pseudobulk UCell score of the individual donor samples (n = 24). Statistical significance was assessed using a one-sided Wilcoxon rank-sum test, with the p-value indicated. **E,** Pseudotemporal ordering of cells of the GC subset visualized as UMAP and violin plots. **F,** Fitted curves showing the expression of representative genes (*STMN2, NEUROD1, NRGN, BHLHE22*, early expression; *RBFOX3, SLC38A11,* late expression) over pseudotime. **G,** Combined expression levels of gene groups defining the ImN transcriptional profile (as detailed in Supplementary Table 3.4) are displayed as UCell scores. Each dot represents the pseudobulk UCell score of the individual donor samples (n = 24). Statistical significance was determined using a one-sided Wilcoxon rank-sum test, with the p-value provided. **H,** Integration of fetal neurogenic lineage and adult GC subset represented as UMAP and as barplot. Arrows indicate the Seurat clusters of integrated datasets where coclustering between adult and fetal cells is observed. See also Supplementary Figure 3J. DiffN, differentiating neurons; HVG, highly variable genes; ImN, immature neurons; MatN, mature neurons; cl, cluster.

### Profiling immature GC transcriptional profiles in the aging human brain

This stringent filtering strategy yielded two major subclusters of mature GC neurons, characterized by the prominent expression of *SGCZ* and *OTOF*, respectively (Figure 3B,C, Supplementary Figure 3D). The *OTOF*-positive population also exhibited high expression of *PDLIM5*, confirming the existence of two main GC subtypes in human DG, as previously reported^17^. Moreover, we identified two subsets of cells expressing neuronal progenitor and differentiation genes (Figure 3B,C, Supplementary Table 3.1), and enriched in expression of previously reported adult human hippocampal immature neuronal signatures^7–9^ (Figure 3D, Supplementary Figure 3E, Supplementary Table 3.3). We annotated these as ‘differentiating neurons’ (DiffN), with one subpopulation specifically expressing *OTOF* (DiffN_OTOF). Within these subpopulations we also observed expression of *ST8SIA2* (Supplementary Figure 3F), a gene encoding for one of the two enzymes that polysialylate the neural cell adhesion molecule (NCAM) into PSA-NCAM, which has been commonly used as histological marker for young neurons in human brain^12–14^. Trajectory inference revealed two *in silico* lineages, both stemming from the DiffN cluster and progressing towards *SGCZ*- or *OTOF*-expressing mature GC (Figure 3E,F, Supplementary Figure 3G,H). We noticed that the DiffN subpopulations were highly abundant (12% of total dataset) and markedly heterogeneous, as suggested by 1) their highly dispersed pseudotemporal distribution in the two lineages (Figure 3E), and, 2) the diverse enrichment for widely-used and previously reported immature neuronal markers within the distinct subclusters (Figure 3G, Supplementary Figure 3E, Supplementary Table 3.3, 3.4). We therefore hypothesized a finer molecular partitioning that could possibly segregate more immature neuronal subsets. To further interrogate the identity of the DiffN clusters and classify with higher confidence the immature cells among them, we integrated the adult GC populations with those in the fetal neurogenic trajectory (Supplementary Figure 3I). We then removed clusters from the integrated dataset that showed no co-clustering between adult and fetal cells. Following this, we used the extent of co-clustering between adult and fetal cells as an indicator of immature neuronal identity. This analysis revealed four clusters in the integrated dataset (Figure 3H, arrows) that consisted of substantial proportions of both fetal and adult cells. Within these, most adult cells were part of the DiffN populations, confirming their transcriptional similarity to fetal cells in the neurogenic trajectory (Figure 3H, Supplementary figure 3J). More specifically, DiffN_OTOF cells and a subset of the DiffN cells (DiffN cluster 13, Figure 3B, Supplementary Figure 3D), showed the highest levels of co-clustering with fetal cells (55.8% and 41.9%, respectively), mainly with NB/ImN (Supplementary Figure 3J,K). Additionally, within DiffN, we identified a subcluster (DiffN cluster 4, Figure 3B, Supplementary Figure 3D) expressing both neuronal differentiation markers (e.g., *STMN1, NRGN, BHLHE22*) and neural stem cell markers (e.g., *CLU, APOE, SOX5*) (Supplementary Table 3.1), with a 15% overlap with fetal progenitors and astrocytes/NSC (Supplementary Figure 3K). These subpopulations co-clustering with fetal cells were also the ones which showed higher enrichment for immature gene sets (Figure 3G, Supplementary Figure 3E), reinforcing their identity as immature neuronal subsets. Subsequently, these cells, comprising DiffN subclusters 4, 13, and DiffN_OTOF, were collectively annotated as immature neurons (ImN) and specifically as ImN_cl4, ImN_cl13, and ImN_OTOF, respectively. In contrast, the remaining subpopulations within DiffN (subclusters 2 and 18) were designated as ImN-like, as they appear distinct from both mature and immature neuronal counterparts (Figure 3I, J). Notably, the proportion of ImN in our dataset reflected the overall enrichment of niche-resident cells compared to previous studies (Supplementary Table 3.5).

To further interrogate their differentiation state, we monitored transcriptional signatures associated with metabolic pathways that are involved in neuronal differentiation. In particular, a metabolic shift from glycolysis to lipogenesis is characteristic of the transition from NSC towards activation and differentiation along the neurogenic trajectory^26,27^. ImN clusters showed significantly higher expression of gene signatures related to glycolysis and decreased levels of lipogenesis genes, further supporting an immature identity (Supplementary Figure 3L).

### Molecular pathways of age and identity in immature neurons

Cellular alterations related to AD- or age-dependent transcriptional programs may lead to a ‘hypo-mature’ or ‘dedifferentiation’ neuronal identity, associated with increased indices of neuroinflammation and neuronal death^28–30^. To further interrogate the nature of the immature neuronal signatures we identified, we compared transcriptional scores of pertinent biological pathways across the adult GC subpopulations (Figure 4A, Supplementary Table 4.1). Interestingly, ImN clusters displayed lower scores of neuroinflammation and neuronal death than the mature ones (Figure 4B, Supplementary Figure 4A), reinforcing the view that the ImN signatures we identified in our dataset are not the result of neuronal dysfunction, like dedifferentiation or fate loss.

**Figure 4.**
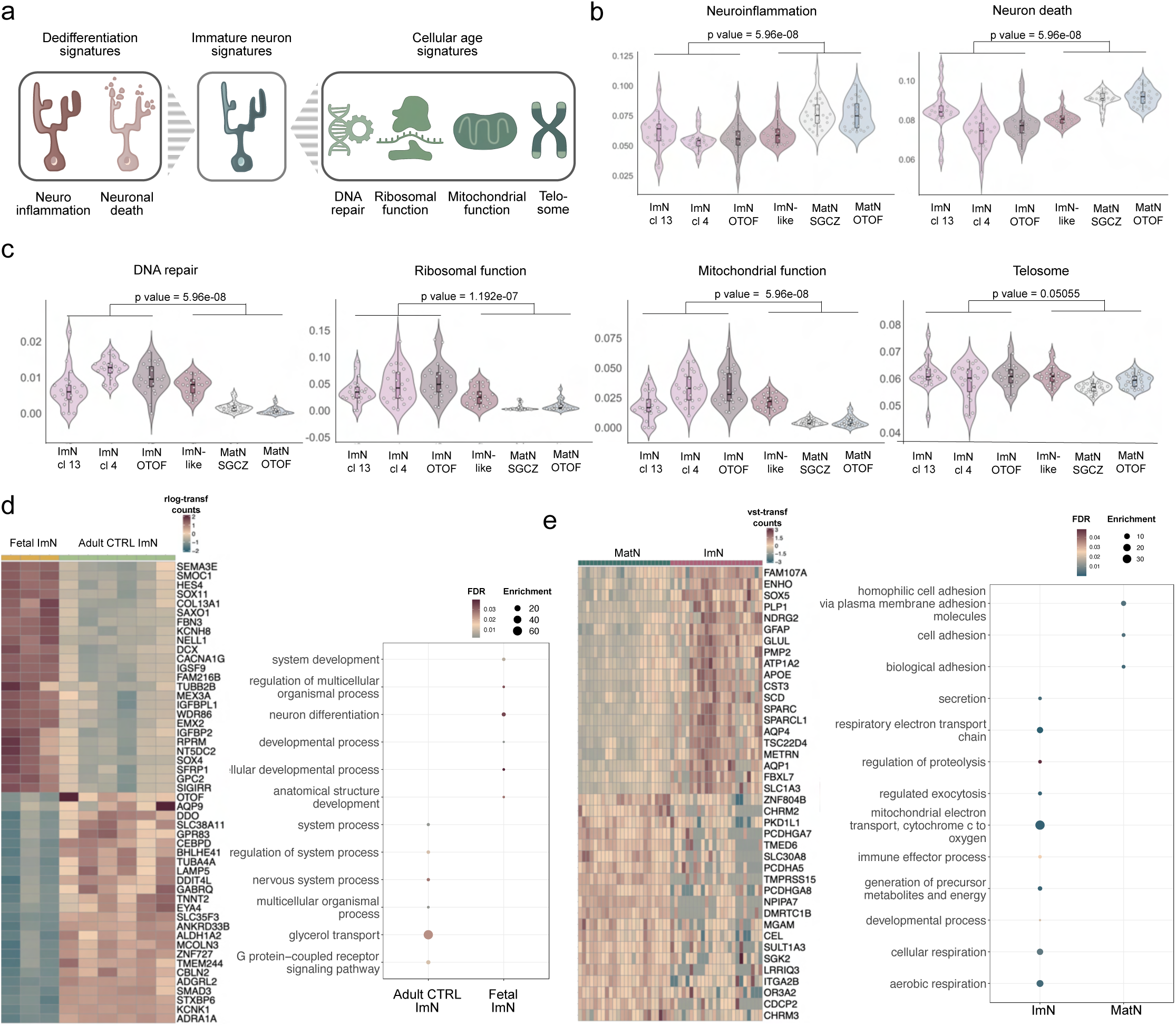
Transcriptional features of immature neurons. **A,** Schematic representation of transcriptional signatures analyzed: dedifferentiation (neuroinflammation and neuronal death) and cellular age-associated pathways (DNA repair, ribosomal function, mitochondrial function, and telosome maintenance). **B, C,** Violin plots showing transcriptional scores for dedifferentiation signatures **(B)** and cellular age-associated pathways **(C)** (Supplementary Table 4.1). Single dots represent pseudobulk UCell score of the individual donor samples (n=24). One-sided Wilcoxon rank-sum test was applied for statistics and p-value is indicated. **D,** Heatmap displaying DEGs between fetal ImN (n=3) and adult ImN populations (n=6, CTRL only), alongside pathway enrichment analysis among these DEGs. **E,** Heatmap illustrating DEGs between ImN and MatN populations, and pathway enrichment analysis among these DEGs. Normalized read counts [calculated using the variance-stabilizing transformation (VST) or regularized log transformation (rlog) method in DESeq2] are shown in the heatmaps. ImN, immature neuron; MatN, mature neuron; DEGs, differentially expressed genes; CTRL, control.

However, it remained uncertain whether these GC subpopulations represent *bona-fide* adult-born ImN or were instead generated earlier-during fetal or juvenile stages- and, due to the neotenic nature of human brain development, exhibit a form of significantly protracted maturation or are developmentally arrested^31–33^. We, therefore, hypothesized that if these cells were indeed born during adulthood, their ‘molecular age’ should differ from the chronological age of the donor, and hence, from the apparent age of the other GC neurons. In an attempt to comparatively ‘birth date’ the cells within the GC population, we implemented a transcriptional proxy of cellular aging, wherein we probed gene signatures of housekeeping cellular functions that were previously shown to be downregulated in human brain with aging, independently of cell type^34^. Interestingly, the ImN population collectively showed the highest expression of genes involved in DNA repair, ribosomal and mitochondrial function (Figure 4C, Supplementary Figure 4B), all previously shown to decrease with age in human brain. In addition, the expression of genes involved in the formation of the telosome, also known as shelterin complex, which is required for the maintenance of the telomere structure and length^35^, was significantly higher in ImN and ImN-like populations compared to the MatN cluster (Figure 4C).

Despite showing some transcriptional similarity, differential expression analysis between the adult ImN pool versus the fetal NB/ImN overall suggested a more pronounced developmental profile for the fetal cells. Conversely, enriched pathways within the adult ImN population were associated with neurotransmission and synaptic plasticity (Figure 4D, Supplementary Table 4.2, 4.3). Among the differentially expressed genes (DEGs), we additionally noted higher expression of genes involved in the regulation of immune response (e.g., *LY86, CD74, CLU, IL6R, BIN1*), inhibition of apoptotic cell death (e.g., *BCL2, BCL2L2, IGF1*), cell survival and neurotrophic signaling (e.g., *FOSB, FGF14, NRG3, BDNF*) in the adult ImN, indicative of a series of putatively functional features specifically acquired within the molecular and cellular complexity of the adult hippocampal niche (Supplementary Table 4.2).

We next asked whether their immature identity was the only biological difference between the adult ImN and the mature GC populations. Differential gene expression analysis between the ImN clusters and the pooled mature GC subsets, apart from an increase in the expression of development-related genes (e.g., *APOE, SOX5, SOX6, BEX1, BEX5, FABP7, BHLHE22, NRGN, STMN1*) and GO terms, also revealed enrichment of pathways and significant upregulation of genes related to proteolysis (*UBB*, *PCOLCE2*), immune response (e.g., *CLU, CST3, CD81, MIF*), mitochondrial respiration (e.g., *COX4I1, COX5B, CYCS, COX7C, COX6A1*), and also increased expression of promyelination genes (e.g. *PLP1*, *PMP2*) (Figure 4E, Supplementary Table 4.4, 4.5).

### Validation of human immature neuronal profiles *in situ* and *in vitro*

To further validate the presence of ImN cells, we next assessed their spatial occurrence in the DG of an independent healthy adult donor (71 years of age, Supplementary Table 1.1) using RNAScope. To select specific markers for ImN, we performed differential expression analysis against both mature GC and astrocytes (Supplementary Table 5.1), thereby identifying genes unique to the ImN population. For mature neuron markers, we performed differential expression analysis between mature neurons and the combined ImN/ImN-like populations to ensure the selection of genes highly specific to mature neurons, that would result in a more distinct expression pattern (Supplementary Table 5.2). For probe selection, we next selected genes whose combined expression was absent from other DG cell types in our dataset (Figure 5A, Supplementary Figure 5A). To improve detection resolution and account for the non-binary expression of ImN markers across GC subsets, we employed a combinatorial expression strategy using probes against multiple transcripts (*NRGN*, *BHLHE22*, *SLC38A11*), allowing us to assess distinct gene expression patterns in individual cells (Figure 5B,C). Using this method, we successfully delineated our four GC subpopulations, namely ImN, ImN_OTOF, MatN_OTOF, MatN_SGCZ, each characterized by distinct combinatorial expression levels of the targeted transcripts (Figure 5D, Supplementary Figure 5B-D, Supplementary Table 5.3). ImN marker labeling was only observed within the GCL and SGZ of the DG, and not elsewhere in the hippocampus. We further verified the identity of the labelled adult cells by labeling for *NRGN*, *BHLHE22* and *SLC38A11* in fetal hippocampus (Figure 5E). There, the *in situ* profile of *NRGN* and *BHLHE22* colocalized with the markers of fetal NB/ImN, *EPHA7* and *SEMA3C*, while *SLC38A11* did not label any cells, given the absence of fully mature neuronal populations at gestational week 24^20,21^.

**Figure 5.**
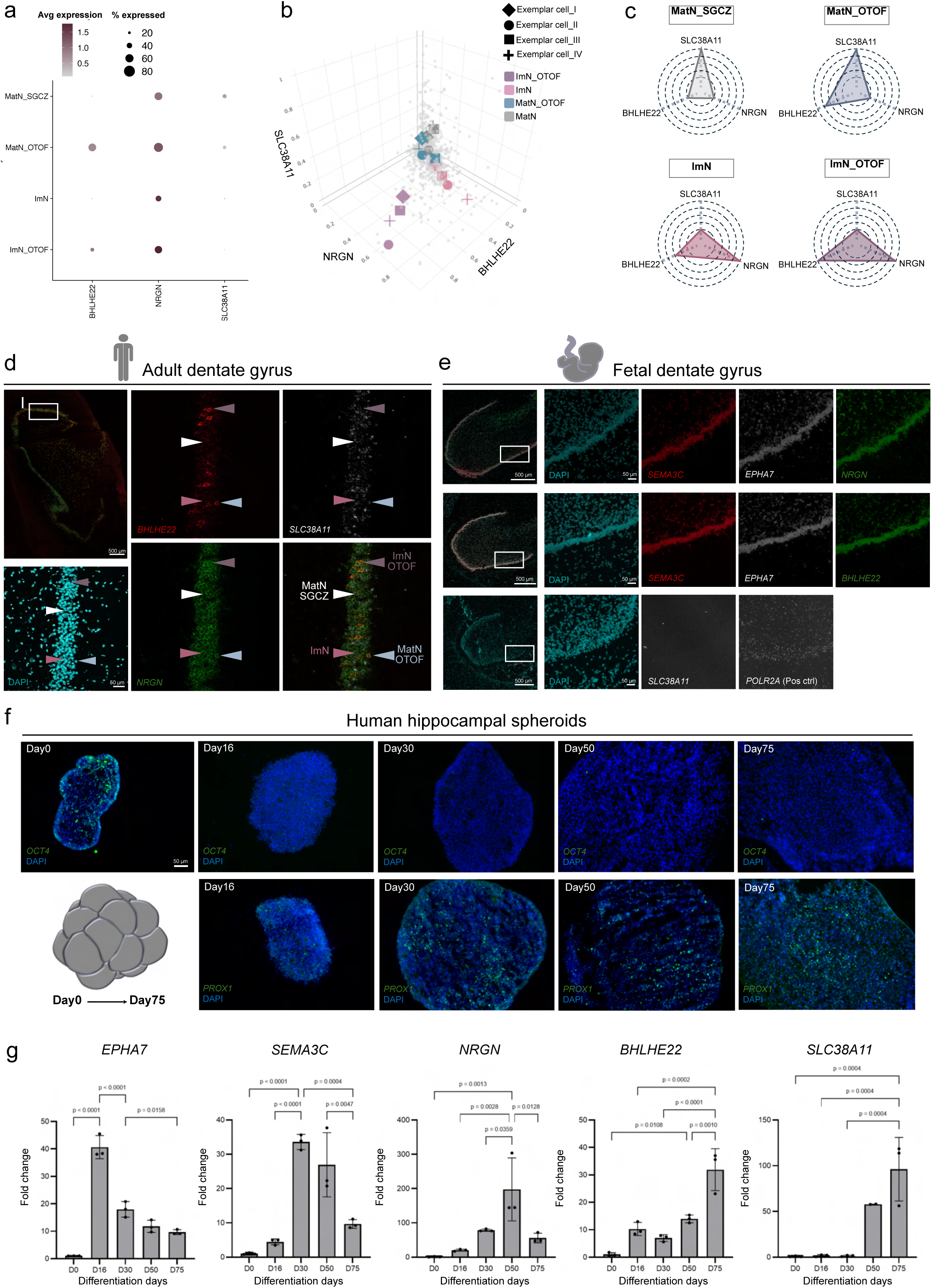
*In situ* and *in vitro* validation. **A,** Dot plot depicting expression profiles of selected marker genes (*NRGN*, *BHLHE22*, and *SLC38A11*) across GC subpopulations. **B,** 3-dimensional visualization of marker gene expression captured via RNAScope *in situ* hybridization. Each dot corresponds to a single cell identified through segmentation analysis of the analyzed images. Exemplary cells representing each GC subpopulation are highlighted for each analyzed image (see also panel D and Supplementary Figure 5B, C). **C,** Spider charts illustrating combinatorial expression patterns of selected genes in the four GC subpopulations. **D,** RNAScope validation of GC subpopulations in an independent healthy adult donor. Arrowheads indicate cells exemplary of each subpopulation. For color code, see B. **E,** Transcript expression pattern measured via RNAScope in the fetal DG (gestational week 24). **F,** Immunostaining of human iPSC-derived hippocampal spheroids over 75 days of differentiation. *OCT4* (Day 0) confirms the pluripotent state, while *PROX1* expression marks differentiation towards GC. **G,** Semi-quantitative qPCR analysis of gene expression in hippocampal spheroids during differentiation. Data are shown as mean ± standard deviation (SD). Statistical comparisons were performed using One-way ANOVA followed by Tukey’s test; p-value is indicated. ImN, immature neuron; MatN, mature neuron.

Considering the congruence of these transcriptional signatures in adult and fetal brain, we next used human induced pluripotent stem cell (iPSC)-derived hippocampal spheroid (three-dimensional) cultures to monitor patterns of gene expression during human neuronal differentiation (Figure 5F, Supplementary Figure 5E-G). *NRGN* and *BHLHE22* became upregulated early along the differentiation trajectory, similar to *EPHA7* and *SEMA3C*, suggesting a role in neuronal differentiation (Figure 5G). *SLC38A11* expression, on the other hand, increased later, which may indicate its importance for neuronal maturation (Figure 5G). Notably, while *EPHA7* and *SEMA3C* showed similar expression profiles during human iPSC differentiation towards cortical neurons (re-analysis from Burke et al.^36^), as those observed in hippocampal spheroids, *NRGN* and *BHLHE22* were expressed at lower levels, and *SLC38A11* expression was nearly undetectable during cortical neuron differentiation (Supplementary Figure 5H). This divergent expression may indicate a specific function for *NRGN*, *BHLHE22* and *SLC38A11* for human hippocampal neurogenesis.

### ImN gene expression programs in Alzheimer’s disease and resilience

We then sought to map distinct ImN gene programs in our four donor groups, namely CTRL, MAD, SAD and RES individuals. No differences in cell proportions were noted across groups (Supplementary Figure 6A). However, we identified significant differences in their transcriptional profiles. The largest number of differentially expressed genes was observed between SAD and RES donors (Figure 6A,B, Supplementary Table 6.1). In SAD ImN, there was also a non-significant but consistent trend of decreased expression in reported ImN transcriptional profiles^7–9^ (Supplementary Figure 6B, Supplementary Table 3.3, 3.4). Notably, the levels of markers of neuronal immaturity *STMN1* and *STMN2*^8,9^, were significantly lower in SAD when compared to RES (Figure 6C). In addition, *PEA15*, a negative regulator of apoptosis^37^, *UBL5*^38^ and *USP11*^39^, involved in DNA repair, and *PSAP*, essential for neurotrophic signaling and neuronal survival^40^, were all significantly lower in SAD when compared to RES. Conversely, the expression of proinflammatory genes, like *PDE4B*^41^*, HECTD2*^42^, *SCART1*^43^, was increased in SAD compared to RES ImN. *NDRG2*, highly expressed in proliferating NPCs and involved in neurogenesis during embryogenesis, postnatal life and upon injury^44,45^, decreased in SAD versus RES donors (ImN cluster 4). Similarly, *DHFR*, which exhibits marked expression in human and mouse NPCs at early stages of neocortical development and was reported to control neurogenic transitions^46^, was also decreased in the SAD group when compared to RES samples (ImN cluster 13). The levels of *ANKFN1*, a gene involved in the regulation of cellular polarity and previously shown to be upregulated in human AD hippocampus^47^, were lower in RES compared to CTRL and MAD ImN. Interestingly, the expression of *CNTN4*, which codes for a protein that interacts with APP to regulate both synaptic plasticity and immune response^48,49^, increased in MAD compared to RES samples (ImN cluster 13). In SAD compared to CTRL ImN, genes related to the suppression of Aβ production and regulation of apoptosis in aging (*APMAP, PDE5A*) were significantly downregulated^50,51^. Interestingly, there were no DEGs between MAD and CTRL in ImN, while DEGs were observed in the comparison between SAD and CTRL ImN, as well as in MAD versus CTRL MatN (Supplementary Table 6.1, 6.2).

**Figure 6.**
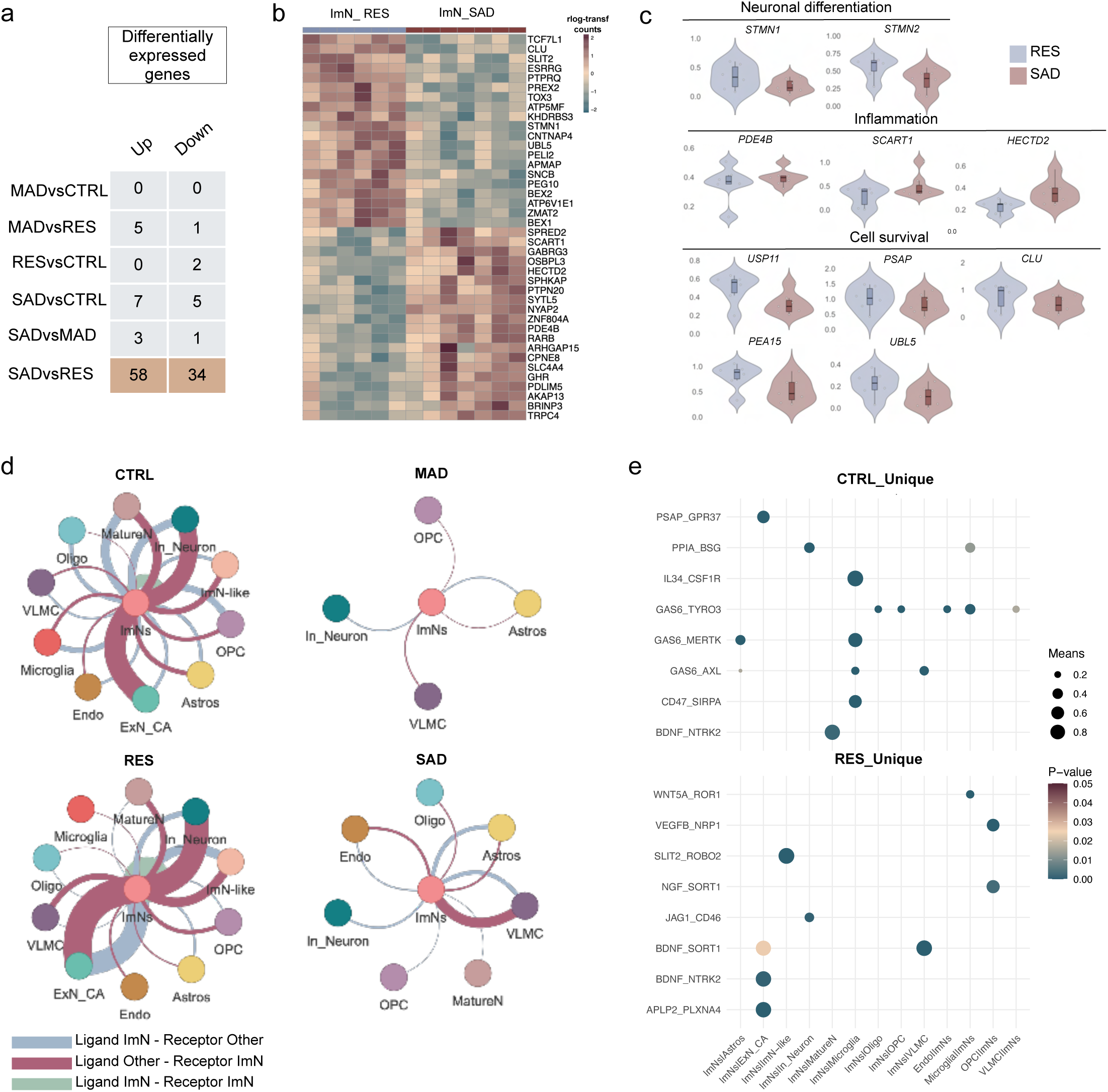
Distinct ImN gene expression programs and intercellular interactions in AD and resilience. **A,** Summary table showing the number of upregulated and downregulated differentially expressed genes (DEGs) in immature neurons (ImN) between the indicated donor groups: MAD (moderate Alzheimer’s Disease), RES (resilient), SAD (severe Alzheimer’s Disease), and CTRL (Control). **B,** Heatmap illustrating top DEGs in ImN, comparing SAD and RES groups. Normalized read counts [calculated using regularized log transformation (rlog) method in DESeq2] are shown. **C,** Violin plots of gene expression for selected DEGs related to inflammation (*PDE4B*, *SCART1*, *HECTD2*), cell survival (*USP11*, *PSAP*, *CLU*), and neuronal differentiation (*STMN1*, *STMN2*) in SAD versus RES ImN. Each dot represents a pseudobulked sample (SAD=7, RES=6). Relative adjusted p-value and fold change are reported in Supplementary Table 6.1, highlighted genes. **D,** Weighted network plots illustrating intercellular signaling of unique interactions centered on ImN in MAD, CTRL, SAD, and RES groups. **E**, Dot plot depicting representative unique ligand-receptor pairs for RES and CTRL.

A particularly interesting observation was that the gene with most consistently higher levels in RES versus SAD in several GC populations, including the pooled ImN cells (Figure 6C), ImN subclusters (ImN_OTOF, ImN cluster 4), ImN-like, and mature subpopulations (Supplementary Table 6.1, 6.2), was *CLU,* coding for a pleiotropic apolipoprotein with antiapoptotic and neuroprotective functions^52,53^. Genetic variation in the CLU-encoding gene has been strongly associated with AD through replicated genome-wide studies (GWAS)^54^, although a functional role in neurogenesis has not been previously reported. Notably, the expression of *CLU* was higher in end-stage differentiating hippocampal neurons (two-dimensional mono-cultures) derived from human iPSCs carrying the APOE2 genotype, which has been shown to confer protection against AD^55^, compared to cultures carrying the APOE4 allele -the most common genetic risk factor for AD- or the APOE3/4 genotype^56^ (Supplementary Figure 6C,D).

To explore variations in cell-cell communication across donor groups, we conducted an *in silico* analysis centered on ImN using CellPhoneDB, focusing specifically on group-unique interactions. This analysis revealed that most of the cellular populations in MAD and SAD exhibited either substantial depletion (InN, oligodendrocytes, OPC, MatureN), or even total absence (ExN_CA, microglia, ImN, ImN-like) of cell type-specific intercellular interactions, compared to CTRL and RES (Figure 6D, Supplementary Table 6.3). In CTRL, several unique ‘signals’ from ImN towards glial populations (oligodendrocytes, OPC, VLMC, astrocytes) included the ligand GAS6 with its receptors TYRO3 and AXL, which are involved in anti-inflammatory, neurotrophic, neuroprotective, (re)myelination, and anti-apoptotic signaling^57,58^. Interestingly, CTRL ImN retained specific interactions with microglia required for physiological phagocytosis of apoptotic cells (GAS6-MERTK, from ImN to microglia)^59^, prevention of phagocytic elimination of healthy cells (CD47-SIRPA, from ImN to microglia)^60^, and general microglial homeostasis and survival (IL34-CSF1R, from ImN to microglia)^61^. Unique interactions between CTRL ImN and mature neuronal populations were also involved in neurotrophic signaling (BDNF-NTRK2, from ImN to MatureN)^62^ and neuroprotection (PSAP-GPR37, from ImN to ExN_CA)^63^ (Figure 6E).

Similarly, ImN in RES would be either the source (BDNF-SORT1, from ImN to VLMC/ExN_CA; BDNF-NTRK2, from ImN to ExN_CA) or the target (NGF-SORT1, from OPC to ImN) of neurotrophic signaling-related interactions^64^ (Figure 6E). Additionally, RES ImN showed unique interactions involved in the regulation of complement activation and immune response (JAG1-CD46, from ImN to InN)^65^ and in non-canonical Wnt signaling that has been associated with tissue repair in ageing and upon injury (WNT5A-ROR1, from microglia to ImN)^66^. A particularly interesting unique ‘signal’ from InN towards ImN in the RES group was that of the APP-SORL1 ligand-receptor pair. SORL1, a sorting receptor for guiding APP to a non-amyloidogenic pathway, is an important AD risk gene, and its brain concentration is inversely correlated with Aβ levels in mouse models and AD patients^67^ (Figure 6E, Supplementary Table 6.4).

## Discussion

Whether or not the human brain can regenerate or repair itself has been a contentious topic for decades. Despite the conflicting views, there is a consensus about conceptual and methodological variables that can confound studies on human adult neurogenesis^4,68–70^. The prospect of *ad libitum* addition of new neurons during aging or in neurodegeneration is undoubtedly attractive from a clinical perspective. Yet, it inevitably raises a host of reasonable questions: Why can’t we profile a ‘complete’ neurogenic trajectory in the human brain, as we have done in rodents^8,9^? What would be the evolutionary advantage of sustained adult neurogenesis in humans^69,70^? What could be the functional relevance of the existence of ‘juvenile’ neurons in neurological conditions?

Here, we established a novel experimental and computational pipeline to reliably identify and profile immature neuronal signatures in aged healthy, AD and resilient human brain. We find significant differences in the transcriptional profiles of immature neurons across groups, suggestive of distinct gene expression programs associated with pathology and cognition.

Using a set of high-quality and short-postmortem delay brain samples, we initially performed a thorough donor stratification to account for potential confounders. To ensure sufficient statistical power^71^, we employed a micropunching-based approach to enrich for the anatomical subfield of the presumed human hippocampal neurogenic niche, and further aimed for high sequencing depth. This allowed us to effectively enrich for the presumed neurogenic niche, achieving a yield of 54% GCs in the total dataset, which is approximately two to seven times more than in other human hippocampal snRNA-seq studies so far^7–9,15–17^. To further filter out from the resulting dataset any RNA contamination-related profiles that could potentially either obscure or artefactually generate neurogenic signatures, we devised a pipeline of rigorous quality control steps not previously implemented in similar datasets. This strategy enabled us to identify two main populations with ImN characteristics: a first one (ImN cluster 13 and 4), marked by high expression of previously identified ImN markers^7–9^, and a second, which additionally expressed high levels of *PDLIM5, OTOF* and *POSTN* (ImN OTOF)^72–74^. Interestingly, a distinct *PDLIM5*^+^ population has not been previously identified in mouse DG^75^, while a *PDLIM5*^+^-cluster has been previously reported as a subset of adult human GC neurons^17^, putatively suggesting a primate-specific identity.

Delineating the full neurogenic trajectory in the adult human brain, from NSC, via proliferating intermediate precursor cells, to further differentiated immature and subsequently mature neurons, would offer compelling evidence for the existence of adult neurogenesis, as we know it from rodent studies. Yet, we and others recently showed that mouse-inferred neurogenic signatures are not suitable for probing adult neurogenesis in humans^4,8,9^. Would one then expect to find the same subsets of transitioning neurogenic populations in the human brain? Interestingly, among the markers of ImN subcluster 4 we noticed several genes highly expressing NSC marker genes. However, this transcriptional profile alone does not conclusively indicate that these cells occupy an earlier stage in the neurogenic trajectory.

Of note, we could not identify any proliferating NPCs in our adult dataset, which at first glance could be indicative of the absence of cell division in the adult human neurogenic niche. Yet, none or minute proportions of adult NPCs have been previously reported not only in human^7,8^, but also in macaque^17^, and even in mouse adult brain^75,76^ using sn/scRNAseq. This may suggest that capturing cycling NPC populations with single-cell transcriptomics, especially in postmortem human brain tissue, may require particularly large sample sizes or marker-based enrichment strategies^73,77^.

The identification of ImN signatures *per se* does not constitute definitive evidence for the existence of adult-born neuronal populations in human brain. Rather, these ImN subsets may represent developmentally born neurons undergoing protracted maturation, as human neurons can take years to fully develop, unlike neurons in rodents, which reach maturity within weeks^32,78^. This phenomenon of human brain neoteny is thought to be a prerequisite for optimizing and increasing the complexity of neural circuits and also for rendering the human brain more susceptible to nurture^79,80^. In our samples, adult ImN displayed transcriptional profiles corresponding to lower levels of neuroinflammation, neuronal death and cellular aging compared to the other GC neurons. Although precise cellular birthdating using transcriptomics data remains challenging^81^, these profiles suggest a more ‘juvenile’ cellular identity^82^. Supporting this, a recent study using adult human brain slice cultures demonstrated that new neurons can be generated in the adult human hippocampus^8^.

Irrespective of when they may have been born, ImN in the adult human brain displayed unique gene signatures and transcriptional pathways that were distinct from both fetal ImN and adult mature neuronal populations. These transcriptional signatures included genes critical for immune response, neurotrophic signaling, neuronal survival and mitochondrial function, hinting that ImN in the adult human hippocampus may not merely represent dormant cells halted within the neurogenic trajectory. Instead, they may have acquired specialized non-canonical roles that support the overall resilience and plasticity of the hippocampal niche microenvironment. This aligns with findings that increased inflammation, mitochondrial dysfunction and loss of proteostasis are hallmarks of cellular aging^83–85^. In the case that the identified adult ImN are developmentally derived rather than generated in adulthood, our findings may suggest an active functional role for neoteny in the adult human brain: preserving a state of developmental plasticity within a distinct cellular subpopulation may be required for tissue homeostasis in the adult human hippocampus.

When comparing ImN across our donor groups, we did not observe differences in cellular composition, as opposed to an earlier snRNA-seq study reporting reduced ImN numbers in AD versus healthy brain^8^. This discrepancy emphasizes the importance of interindividual variability, as a substantial subset of control donors in the earlier study did not exhibit a higher proportion of ImN relative to AD donors. Differences in technological platforms, wet and dry lab pipelines, or sample size, may all account for this deviation, as well. Yet, we did identify unique group-specific gene expression signatures. Of note, the largest number of DEGs was found when comparing SAD to RES samples, which could suggest that transcriptional, and perhaps also functional programs in these two groups exhibit maximal divergence. SAD ImN differed from RES ImN in gene signatures related to neurogenesis, inflammation, cell death and apoptosis. Interestingly, when comparing MAD to CTRL, ImN did not exhibit any transcriptional alterations, as opposed to what was observed in the SAD versus CTRL comparison. In addition, MatN did harbor significant DEGs already in MAD compared to CTRL. These observations could suggest that ImN are less severely impacted at initial stages of pathology progression. The DG is relatively spared until mid-late AD compared to other brain regions^86^. A multivariate stressor-threshold theory suggests that chronic accumulation of disruptions in cellular homeostasis may underlie cellular vulnerability in neurodegeneration^87^. ImN may actively enhance homeostasis in the niche microenvironment, and thereby contribute to the observed ‘fitness’ of the DG in the beginning of pathology progression, although replication studies and larger sample sizes are needed to further support this hypothesis.

An overall decrease in unique interactions between ImN and other niche-resident cellular populations was observed in MAD and SAD compared to CTRL and RES, indicating a putative deficiency in intercellular crosstalk with AD pathology and cognitive decline. Similar changes were previously documented for ImN in AD patient brain^8^, while an overall decline in intercellular communication in human AD prefrontal cortex has also been previously reported^88,89^. Interactions between ImN and other cell types in CTRL samples were involved-among others- in anti-inflammatory, anti-apoptotic, neurotrophic, and neuroprotective signaling, suggesting that ImN may play key roles in maintaining niche homeostasis under physiological conditions.

In RES samples, ImN additionally contributed to cell-cell interactions involved in the regulation of immune response and tissue repair, indicative of possible resilience-related modulatory roles. Reduced inflammation emerged as a common outcome across various geroprotective interventions in mice, including senolysis, caloric restriction, *in vivo* partial reprogramming, and heterochronic parabiosis^90^. In human brain, lower levels of inflammation are associated with resilience to AD pathology^91–93^. Interestingly, a unique interaction between InN and ImN in the RES group was that of the APP-SORL1 ligand-receptor pair. *SORL1* variants are associated with AD risk, while SORL1 reduces β amyloid levels in the brain by preventing the amyloidogenic processing of APP and also inducing lysosomal degradation of β amyloid^94^. This could indicate a resilience-specific activation of an anti-amyloid mechanism within ImN in the presence of AD pathology. Overall, these findings suggest that ImN play important roles as a component of the multicellular hippocampal network in healthy, AD and resilient human brain.

Our work has several limitations. First, transcriptional profiling in postmortem human brain only provides a snapshot of gene expression and, alone, cannot infer functional output of the identified cells or accurately depict intercellular and lineage relationships among them. Second, the strategy we employed to identify ImN signatures may have overlooked certain potentially relevant populations, particularly those lacking transcriptional similarities to fetal ImN. Third, our analysis across the four distinct donor groups focused on ImN, and did not explore the potential contribution of other cellular populations to mechanisms underlying AD pathology and resilience. Fourth, we cannot exclude that our resilient donors might have progressed towards becoming cognitively impaired if they were to live longer. Nevertheless, when compared to age-matched AD patients with comparable levels of amyloidosis, they demonstrated better cognitive performance, suggesting distinct responses to pathology. Fifth, the two- and three-dimensional hippocampal neuronal cultures that we used are of embryonic-like nature, and may, thus, only provide partial representation of the neuronal differentiation process occurring in the adult human brain.

In conclusion, our study identifies unique transcriptional signatures in adult human ImN that may play pivotal roles in healthy and resilient brain. Regardless of whether these represent *de novo* adult immature or developmentally born cellular populations, understanding their functional roles in the aged, degenerating and resilient brain is crucial. Our dataset can be used as a resource for identifying genes and gene programs with potential therapeutic relevance, offering a foundation for mapping novel targets in the treatment of AD.

## Methods

### Human brain tissue

Fresh frozen hippocampal tissue was obtained from the Netherlands Brain Bank (NBB) from 24 adult donors. Informed consent for brain autopsy and the use of brain tissue and associated clinical data for research was obtained by the NBB in accordance with international ethical guidelines. Autopsy procedures were conducted following approval from the Medical Ethics Committee of Vrije University Medical Center in Amsterdam, the Netherlands. Donors were selected based on comprehensive neuropathological assessment, clinical data, and either Clinical Dementia Rating (CDR) or Global Deterioration Scale (GDS) scores. Neuropathological assessment followed standardized protocols, including Braak staging^95^, CERAD guidelines^96^, and the National Institute on Aging– Alzheimer’s Association criteria^97^. Cases with psychiatric or neurological disorders unrelated to AD, or with severe comorbid pathologies such as cortical Lewy bodies, hippocampal sclerosis, or LATE (Limbic-predominant age-related TDP-43 encephalopathy), were excluded. CDR or GDS scores were used retrospectively to assess cognitive function within the three months prior to death.

The cohort included individuals with Alzheimer’s disease (AD) and advanced dementia (CDR 2-3) with high to intermediate AD pathology (Braak stages 4–6, Thal phase ≥3), resilient donors with preserved cognition (CDR 0–0.5) despite intermediate to high AD pathology, and cognitively intact, age-matched controls with minimal AD pathology (Braak stages 0–2, Thal phase ≤2). Donors were matched for sex, age, postmortem interval, brain pH, and APOE genotype. Information about donor cohort is summarized in Supplementary Table 1.1. Fresh frozen hippocampal tissue was obtained from the Amsterdam University Medical Center (UMC) and the Dutch Fetal Biobank from 3 aborted fetuses (gestational week 24), in accordance with the Informed Consent Standards for Human Fetal Tissue Donation and Research developed by the International Society for Stem Cell Research.

### DNA isolation and genotyping

DNA was extracted from 50 mg of cerebellar tissue using the DNeasy Blood & Tissue Kit (Qiagen, Netherlands). The tissue was lysed overnight at 56°C, after which the lysate was passed through DNeasy Mini Spin Columns (Qiagen, Netherlands) and processed following the manufacturer’s protocol. The Apolipoprotein E (ApoE) genotype was determined using the TIB Molbiol LightMix Kit APOE C112R R158C (Roche Diagnostics, Netherlands), utilizing the LightCycler® 480 System with the hybridization probe method.

### RNA isolation and RNA integrity number (RIN) assessment

A tissue sample of 15–20 mg derived from trimmed DG and cortex was used for RNA isolation using the RNeasy Mini Kit (Qiagen, Netherlands) combined with Trizol (3 ml Trizol per 100 mg of tissue; Thermo Fisher Scientific, Netherlands). Following homogenization, phase separation was carried out by adding chloroform, followed by vigorous shaking and incubation at room temperature (RT) for 2–3 min. The samples were then centrifuged at 12,000 g for 15 min at 4°C. The aqueous phase was mixed with an equal volume of 70% ethanol. Samples were then loaded on a RNAeasy Mini column (Qiagen, Netherlands) and further processed according to the manufacturer’s instructions. RNA yield and purity was determined using Nanodrop (Thermo Fisher Scientific, Netherlands). Quality of RNA was determined with 4200 Tapestation (Agilent Technologies).

### Single-nucleus RNA sequencing

#### Nuclei isolation from dentate gyrus

To enrich for specific cell types within the subgranular zone (SGZ) and granular cell layer (GCL) of the dentate gyrus, fresh-frozen adult human hippocampal tissue blocks were sectioned into 300 μm thick slices using a cryostat (Leica, Netherlands). Sections were maintained on dry ice, and SGZ and GCL, extending from the molecular layer to the hilus, were isolated under a stereomicroscope using a 0.5 mm-diameter biopsy puncher. Adjacent sections were stained with hematoxylin and eosin to facilitate the accurate identification of the dentate gyrus. Punched tissue samples were stored at −80°C until further processing. For the fetal samples, the region of the tissue containing the DG was identified by hematoxylin and eosin staining and sectioned into 50 μm-thick slices using a cryostat (Leica, Netherlands).

Nuclei were isolated from the frozen punched tissue or sections using an optimized isolation protocol^98^. Briefly, samples were homogenized using a glass Dounce homogenizer (15 gentle strokes) in 1 mL of ice-cold Homogenization Buffer (HB), which contained 320 mM sucrose, 5 mM calcium chloride, 3 mM magnesium acetate, 10 mM Tris, 0.1 mM EDTA, 0.1% Igepal, 0.1 mM PMSF, 1 mM 2-mercaptoethanol, and UltraPure water, along with 5 μL RNasin Plus (Promega, Benelux, Belgium). The homogenate was filtered using a 70 μm strainer and washed with 1.65 mL HB, bringing the final volume to 2.65 mL. An additional 2.65 mL of Gradient Medium (containing 5 mM calcium chloride, 50% OptiPrep (Sigma-Aldrich, Belgium), 3 mM magnesium acetate, 10 mM Tris, 0.2 mM PMSF, and 1 mM 2-mercaptoethanol) was added to achieve a final volume of 5.3 mL. To isolate the nuclei, OptiPrep Diluent Medium (ODM) consisting of 150 mM potassium chloride, 30 mM magnesium chloride, 60 mM Tris, and 250 mM sucrose was prepared. The sample was layered onto a 4 mL 29% cushion (29% Optiprep, ODM) using a P1000 pipette, and the weight was balanced with HB. Centrifugation was carried out at 7,700 rpm for 30 min at 4°C using a SW41Ti rotor. After centrifugation, the supernatant was removed with a plastic Pasteur pipette, and the lower supernatant was aspirated using a P200 pipette. The nuclei were resuspended in 200 μL of resuspension buffer (1x PBS, 1% BSA, and 0.2 U/μL RNasin Plus), transferred to a fresh tube, and washed again with 100-200 μL of the same buffer. The resuspended solution was pooled, and any clumps were disrupted by pipetting with a P200. The sample was then filtered through a 0.35 μm strainer using a Falcon tube. A 9 μL-aliquot of the sample was mixed with 1 μL of propidium iodide (PI) stain, loaded onto a LUNA-FL slide, and left to settle for 30 seconds. The nuclei were examined under a LUNA-FL Automated Cell Counter to assess numbers and morphology.

#### Single-nucleus microfluidic capture and cDNA synthesis

The nuclei samples were further processed for GEM generation, with targeted nuclei recovery of 10000 nuclei/sample by using 10x Genomics Single Cell 3’ Reagent Kit (v3.1) according to manufacturer’s protocols (10x Genomics, CG000204_Rev_A).

#### Library preparation and sequencing

Post cDNA amplification cleanup and construction of sample-indexed libraries and their amplification followed manufacturer’s directions (10x Genomics, CG000183_Rev_A), with the amplification step directly dependent on the quantity of input cDNA. In order to reach sequencing saturation of ∼65% (∼45,000 reads/nucleus), single nucleus libraries were run using paired end sequencing with single indexing on the NovaSeq6000 platform, following manufacturer’s instructions (10x Genomics, CG000204_Rev_A). To avoid lane bias, multiple uniquely indexed samples were mixed and distributed over several lanes. Supplementary Tables 1.2 and 2.1 provide sequencing information per sample.

### Analysis of single-nucleus RNA sequencing datasets (adult and fetal)

#### Alignment and software

Paired-end FASTQ files were aligned and quantified with STARsolo^99^ from STAR v2.7.10b_alpha_230301(--soloFeatures Gene GeneFull_Ex50pAS Velocyto, --soloType CB_UMI_Simple, --soloCBwhitelist 3M-february-2018.txt, --soloUMIlen 12, --soloCellFilter EmptyDrops_CR, --soloMultiMappers EM) using the human reference genome GRCh38 (GENCODE v32/Ensembl98). All downstream steps were performed using the count matrix of GeneFull_Ex50pAS option, unless specified otherwise.

#### Quality control

After STARsolo quantification, data were processed using CellBender (v. 0.2.2) with the ‘remove-background’ command^25^, which was employed to eliminate systematic biases and background noise caused by ambient RNA and random barcode swapping. The output count matrix from CellBender was imported and further processed by using Seurat (v. 4.4.0) package^100^ in R (v4.3.1). The removal of potential doublets was performed by using scDblFinder^101^ with an expected doublet ratio of 0.7 or higher. For each specimen, genes expressed in <10 nuclei were discarded. To rule out low quality nuclei, the count matrix of each sample was filtered to exclude nuclei with >10% mitochondrial content and < 25% intronic ratio. Intronic ratio was calculated as unspliced counts/(unspliced+spliced counts). For every specimen, a minimum cutoff for number of genes was determined by visual inspection, and samples were filtered accordingly (Supplementary Figure 1B).

#### Normalization, integration and main cell type identification

Following quality control, all nuclei were retained for dimensionality reduction and unsupervised clustering, which were also performed using Seurat. The 24 adult libraries were merged together in a unique dataset and the same process was applied to the 3 fetal libraries. For each merged dataset (adult and fetal), raw UMI counts were normalized using the ‘NormalizeData’ function, with the scaling factor equal to 10,000. Top 2000 variable features were selected with the function FindVariableFeatures, and data were scaled using the *ScaleData* function. The dimensionality of the data was first reduced by principal component analysis (PCA). To account for large differences in expression across patients, as well as for technical batch effects, principal components were aligned with Harmony (v1.1.0 with kmeans_init_nstart=20, kmeans_init_iter_max=150, lambda=1 options)^102^ using ‘sample and ‘run’ as batch indicators. The corrected components (20 for adult, 20 for fetal) where then used for constructing a neighborhood graph and for clustering the nuclei based on transcription similarity using the Louvain algorithm (*FindNeighbors FindClusters*), and applying appropriate resolution (1.4 for adult, 2 for fetal). Finally, the nuclei clusters were visualized with uniform manifold approximation and projection (UMAP). All of the above functions are part of the Seurat R Package 4.4.0. For the adult dataset, clusters with high *RBFOX3* expression were identified as neurons, which were further classified into excitatory or inhibitory subtypes based on the presence of *SLC17A7* or *GAD2*, respectively. Excitatory neurons were divided into granule cells (GC), marked by *PROX1*, and mossy cell/Cornu Ammonis (MC/CA) neurons, distinguished by *CFAP299*. Within the GC population, two subtypes were identified: *SGCZ*- and *OTOF*-expressing cells. The remaining nuclei were annotated as non-neuronal cell types and categorized into astrocytes (*AQP4*, *GJA1*), endothelial cells (*CLDN5*, *CD34*), microglia (*PTPRC*, *CSF1R*), oligodendrocytes (*MOBP*, *MOG*), oligodendrocyte progenitor cells (*PDGFRA*, *NEU4*), and vascular and leptomeningeal cells (*DCN*, *COL1A2*).

Annotation for the fetal dataset followed a similar approach, using markers identified in a previous study of human hippocampal development^103^.

#### Additional quality control steps on GC subset

To discern finer distinctions within the GC population, nuclei annotated as GC were isolated for further analysis. To minimize potential misannotations arising from technical artifacts— such as residual ambient RNA contamination not fully removed by Cellbender or variability in data quality— additional filtering and validation steps were undertaken to enhance accuracy. First, astrocytes and GC were subsetted and re-clustered together to verify that the GC population exhibited a distinct transcriptional profile. Additionally, we integrated the GC population with empty droplets (described below) and removed GC nuclei that co-clustered with empty droplets (n = 1,386 nuclei). To profile ambient RNA in the dataset, differential gene expression analysis was performed between empty droplets and true cells of GC population (pseudobulking data at the sample level, DESeq2). The transcriptomic profile of empty droplets was then removed from top 2000 high variable genes of GC population to ensure clustering is not affected by ambient RNA profile.

Additionally, the GC population was re-clustered by using only the unspliced RNA matrix from Velocyto to further confirm that the clustering is not driven by cytoplasmic (spliced) RNA debris.

#### Identification of empty droplets

To select empty droplets for quality control, a barcode rank versus total UMI plot was generated for each sample, displaying UMI counts per droplet ranked from highest to lowest. True nuclei were identified as all droplets up to the first plateau in the curve. An equal number of empty droplets was then selected from the region surrounding the second plateau, corresponding to empty/ambient RNA-containing droplets (low-count UMI). Specifically, half of the empty droplets were sampled from barcodes ranked just below the total droplet inclusion threshold, and the other half were taken from subsequent barcodes.

#### Fetal-adult integration

The adult GC dataset was integrated with the fetal neurogenic dataset using Harmony^102^ (version 1.1.0) with the following parameters: kmeans_init_nstart=20, kmeans_init_iter_max=150, and lambda=1. Batch effects were addressed using ‘sample’ and ‘agegroup’ as batch covariates. Following integration, clusters that showed no co-clustering between fetal and adult cells were excluded (primarily composed of adult mature GC). The integration and clustering steps were then repeated to refine the analysis.

#### Differentially expressed gene (DEG) computation

To avoid false discoveries^104^, pseudobulk differential gene expression analysis was performed using the DESeq2^105^ (v1.42.0) package in R. The raw gene expression matrix for individual cells was extracted from the counts slot of the RNA assay. Counts were then summed per gene for each sample to create a pseudobulk expression matrix. The resulting matrix was normalized in DESeq2 by estimating size factors and performing normalization through the counts function, resulting in a matrix of normalized counts per gene and sample, analogous to the output of a traditional bulk sequencing experiment. Differential gene expression was calculated using the DESeq function, which fits a negative binomial distribution to the data. To identify markers of cell (sub)type, pairwise comparisons were performed between the cluster of interest and all the remaining clusters without applying any batch correction (design = ∼cluster). When identifying DEGs across disease groups, batch effects due to sex and sequencing run were accounted for (design = ∼0+Sex+Run+group). For group comparisons within a specific cluster, an artificial run batch was created to group single-sample-runs together. Comparisons between fetal and adult clusters were conducted without batch correction. DEGs were filtered based on log fold change (logFC) and adjusted p-values for downstream analyses, as detailed in the respective Supplementary Table legends. DEGs were visualized in heatmaps using rlog-transformed counts for datasets with fewer than 20 samples per condition and vst-transformed counts for datasets with more than 20 samples per condition, as described in the figure legends. Samples from all four groups (n = 24) were used for differential gene expression analysis unless otherwise specified.

#### Gene set score analysis

Gene set scores (Figures 2,3,4, Supplementary Figures 3,4,6) were calculated with the function AddModuleScore_UCell() of UCell R package (v2.6.2) using the gene sets reported in Supplementary Tables 2.6, 3.3, 3.4, 4.1. Samples from all four groups (n = 24) were used.

#### Trajectory inference

In the fetal neurogenic subset and in the adult GC subset, the Slingshot R package (2.10.0) was used to order cells along a pseudotime lineage and a negative binomial generalized additive model was fitted using the *fitGAM* function of the tradeSeq R package (v1.160)^106,107^. Specifically, associationTest() and startVsEndTest() functions were used to determine gene expression changing between start and end of the pseudotime lineage and results were filtered by pvalue<0.05 and abs(logFClineage1)>1. Selected genes were plotted on heatmap using data slot of the RNA assay retrieved by GetAssayData() function of Seurat.

The pseudotime analysis for the adult GC subset was also replicated and compared using the monocle3 R package^108^.

Additionaly, RNA velocity was used in the GC subset to infer transcriptional dynamics and predict the future state and transition of individual nuclei in the dataset using the spliced and unspliced counts data generated by velocyto option of STARsolo. RNA velocity was calculated and visualized by RunSCVELO() function of SCP R package (v0.5.6) ^109^.

#### Compositional analysis

Variations in sample level cell type proportions followed by data transformation and statistical testing across disease groups were calculated with propeller function from speckle R package (v1.2.0)^110^ using “logit” transformation parameter. For statistical testing, F-test (ANOVA) was used, and Benjamini and Hochberg false discovery rates were calculated to account to multiple testing of clusters.

#### Cell-cell interaction analysis

Cell–cell interactions were predicted by CellPhoneDB Python package (v5.0.0)^111^. First, the normalized gene counts matrix and metadata of each cell from each disease group were extracted, and then the statistical analysis method of CellphoneDB was used to analyze cell–cell interactions. Briefly, for each group and cell type, the mean expression of each ligand-receptor interaction pair was calculated. A null distribution was created by shuffling cell type labels 1,000 times and recalculating the mean expression for each interaction pair during each iteration. The significance of each interaction was determined by comparing the observed mean expression to the null distribution, with statistically significant interactions identified using a p-value threshold of < 0.05. Subsequently, all interactions involving cell types were identified for each group, and those centered on ImN cells were extracted. Unique interactions were then determined for each cell type pair within each group, providing insight into cell type-specific communication patterns across conditions. Plots were done with ktplots (v2.3.0) R package.

#### Pathway enrichment analysis

Gene ontology (GO) enrichment analysis was performed using GOrilla^112^. The top 150 differentially expressed genes (DEGs) were used as input list (when DEGs <150 all the DEGs were used), while all genes expressed in at least 0.1% of the dataset that was used to compute DEGs, served as the background list. GO Biological Process Direct pathways were ranked by significance, with pathways exhibiting an FDR below 0.05 selected for further analysis.

### *In situ* hybridization (RNAScope)

To detect single mRNA molecules, RNAScope was performed on frozen adult control postmortem (female, 71 years old, Supplementary Table 1.1) and fetal (GW24) hippocampal slices. 20 μm sections were cut from frozen biopsies and mounted on superfrost plus glass slides. One positive (Homo sapiens *PPIB*), one negative (*Escherichia coli DapB*) control probe and 5 different probes against genes (i.e. transcripts) of interest were used (*BHLHE22*, *NRGN*, *SLC38A11*, *EPHA7*, *SEMA3C*). *In situ* hybridization (ISH) was performed according to the protocol of the RNAScope Multiplex Fluorescent Reagent Kit v2 (ACD, Netherlands; Cat. No. 323100) and RNAScope Ancillary kit for Multiplex v2 (Cat. No. 323120).

Briefly, slides from frozen brains were incubated in cold 4% PFA in 1X PBS for 1 hour at 4°C. Slides were then washed 2 times with 1X PBS and dehydrated in 50%, 70%, 100% and 100% ethanol for 5□min each at RT. After drying the slides for 5□min at RT, H_2_O_2_ was added for 10□min at RT. For antigen accessibility, slides were treated with Protease IV for 20□min at RT. Before applying the probes, slides were washed again twice with water. C2, C3, C4 probes were diluted in C1 probe or probe diluent at a 1:50 ratio and incubated on the slides for 2 h at 40°C. C1 probes were detected with TSA-Vivid Fluorophore 650 or 570 for adult and fetal slides respectively, C2 probes with TSA-Vivid Fluorophore 520, C3 probes with TSA-Vivid Fluorophore 570 or 520 for adult and fetal slides respectively, C4 probes with TSA-Vivid Fluorophore 650, dilution 1:1500. Before mounting, DAPI was added for 30 s at RT to label the nuclei. To help protect fluorescent dyes, ProLong Gold Antifade was added to the slides and coverslips were applied. Slides were dried overnight in the dark at RT and imaged in the following 72 hours with a Leica TCS SP8 confocal system as *z*-stacks using a 20X objective.

### RNAScope image processing and data analysis

Acquired images were analyzed using FIJI^113^ and QuPath^114^. Color deconvolution was applied to separate specific excitation channels for the detection of nuclear counterstains (DAPI) and mRNA probes. Analyses were restricted to nuclei located within the dentate gyrus (DG), ensuring a focused examination of this region.

Nuclear segmentation and contour detection employed a deep-learning model optimized for the DAPI counterstain, implemented using the StarDist extension^115^. Segmentation quality was refined through manual inspection, leveraging supervised cell classification features such as nuclear size, roundness, and detection probability to maximize accuracy. X-Y coordinates of cell nuclei were recorded during data collection.

Probe signals were quantified as intensity values within each nucleus or cell and normalized to account for the respective nucleus/cell area. For comparative analyses across different images, probe signal measurements were further normalized to a scale of 0 to 1.

Cell type annotation was performed using a combination of quantitative probe signal measurements and visual inspection, ensuring robust classification within the dataset. QuPath annotation files containing all measurements are available upon request and exported measurement data is provided in Supplementary Table 5.3.

### Human iPSC-derived three-dimensional hippocampal spheroid cultures

Human iPSCs were derived by reprogramming fibroblasts from a healthy donor as previously described^116^. Hippocampal spheroids were subsequently generated using established methods^117^. Briefly, iPSCs colonies were dissociated with dispase and transferred into ultra-low attachment flasks (Corning, USA) containing Wicell medium (DMEM/F12, 20% KSR, 1% NEAA (v/v), 1% L-Glutamine (v/v), and 50 mM β-mercaptoethanol) supplemented with 20 ng/mL fibroblast growth factor 2 and 20 μM ROCK inhibitor Y-27632 (Selleck Chemicals, USA). After 24 hours, the medium was replaced with neural induction medium consisting of advanced DMEM/F12, 2% B27 Supplement without vitamin A (v/v), 1% N2 Supplement (v/v), 1% non-essential amino acids (NEAA, v/v), 2 mM L-glutamine, and 1% penicillin-streptomycin (v/v). For dorsomedial telencephalic neural specification, the medium was enriched with LDN-193189 (Stemgent, USA, 0.1 μM), Cyclopamine (Selleck Chemicals, USA, 1 μM), SB431542 (Sigma-Aldrich, USA, 10 μM), and XAV-939 (Tocris Bioscience, USA, 5 μM) during the first 10 days, with medium changes every other day. On day 10, the free-floating spheres were moved to neuronal differentiation medium (NDM) containing Neurobasal medium, 1% N2 (v/v), 1% NEAA (v/v), L-glutamine, and 1% penicillin-streptomycin (v/v). To promote hippocampal differentiation, the NDM was supplemented with 0.5 μM CHIR-99021 (Stemgent, USA) and 20 ng/mL brain-derived neurotrophic factor (BDNF, PeproTech, USA) for 65 days, with medium changes every other day. Samples were collected at timepoints D0, D16, D30, D50, D75 for semi-quantitative real-time PCR and immunostaining.

### Human iPSC-derived two-dimensional hippocampal neuronal cultures

Gene expression analysis in human iPSC-derived hippocampal neuronal cultures was performed on RNA generated in a previous study^118^. Briefly, the following human iPSCs were obtained from the European Bank for induced pluripotent Stem Cells (EBiSC): ApoE3/E4 (C, original iPSC line), ApoE4/- (C4, CRISPR line), ApoE2/- (C6, CRISPR line). Maintenance passaging of iPSCs was done with Versene-ethylenediaminetetraacetic acid (EDTA) solution (Lonza, UK) according to the manufacturer’s instructions. iPSCs were replated 2 or 1 day(s) prior to the initiation of directed differentiation in StemFlex^TM^ medium and after 7□days of neural induction in N2B27□+□4i medium [N2B27 medium: 1:1 mixture of N-2 medium (DMEM/F12 supplemented with 1X GlutaMAX™ (Thermo Fisher Scientific, UK)) and 1X N-2 (Thermo Fisher Scientific, UK) and B-27 medium (Neurobasal medium, Thermo Fisher Scientific, UK) supplemented with 1X GlutaMAX™ and 1X B-27 minus vitamin A (Thermo Fisher Scientific, UK); 4i: 2 μM XAV939 (Sigma Aldrich, UK, Wnt/β-catenin signaling pathway inhibitor), 10 μM SB431542 (Sigma Aldrich, TGF-β signaling pathway inhibitor), 100□nM LDN193189 (Sigma Aldrich, UK, BMP signaling pathway inhibitor), and 1 μM Cyclopamine (LC Laboratories, USA, Sonic hedgehog signaling pathway inhibitor)]. NPCs were expanded until day 21/22 in N2B27 medium. For GC-like neuronal differentiation, NPCs were terminally plated and induced to differentiation in ND medium, consisting of B-27 medium containing 1 μg/ml laminin, 200−nM ascorbic acid (Sigma Aldrich, UK), 1□mM dibutyryl cyclic adenosine monophosphate (Sigma Aldrich, UK), 20□ng/mL brain-derived neurotrophic factor (BDNF) (Peprotech, UK), and 20□ng/ml WNT3A (R&D systems, UK) for the first 7□days (until D28/29), and subsequently in ND medium without Wnt3a until Day 42/43.

Cells were harvested at timepoints D0, D3, D7, D10, D12, D18, D21, D28, D35, D42 for RNA extraction.

### RNA extraction, reverse transcription and semi-quantitative real-time PCR

Human iPSC-derived hippocampal spheroid samples were collected at the indicated time points, rinsed with PBS and snap frozen in liquid nitrogen. RNA isolation from hippocampal spheroids was performed using the miRNeasy Micro Kit (Qiagen, Netherlands) following manufacturer’s instructions. Briefly, the cell pellets were homogenized in QIAzol Lysis Reagent, followed by a 5 min incubation in chloroform (Thermo Fisher Scientific, Netherlands) at RT. After 15 min of centrifugation at 12,000 g at 4°C, the upper aqueous phase was collected, supplemented with 1.5 volumes of 100% ethanol and loaded on RNeasy MinElute spin columns. Following washing steps, RNA was eluted in 14 µl of RNase-free water and measured using the NanoDrop ND-1000 SpeCTRLophotometer (NanoDrop Technologies, Inc.).

Human iPSC-derived hippocampal neuronal cultures were collected at the indicated time point, and total RNA was extracted using the TRIzol® reagent (Thermo Fisher Scientific, UK) according to manufacturer’s instructions.

Reverse transcription of mRNAs for both hippocampal spheroids and hippocampal neurons was performed with 50-200 ng RNA to synthesize cDNA using Superscript II reverse transcriptase (Thermo Fisher Scientific, Netherlands).

Real-time semi-quantitative PCR was performed using the PowerUp™ SYBR™ Green Master Mix (Applied Biosystems™). *GAPDH* and *PSMB4* were used as housekeeping genes. Primer sequences can be found in Supplementary Methods. Ct values were determined using the second derivative method and subsequently fold changes were calculated using the ΔΔCt method.

### Immunostaining, microscopy and image analysis

For immunocytochemistry, cells were washed three times with DPBS, and fixed with 4% PFA for 40 min at 4°C. After three additional washes with DPBS, cells were permeabilized with 0.3% Triton in PBS for 30 min at RT, and blocked for 1 hour at RT in 10% normal donkey serum (NDS, VWR, USA) in PBS containing 0.1% Triton-X (Sigma-Aldrich, USA). Cells were incubated in target-specific primary antibodies overnight at 4°C in 10% NDS in PBST. Subsequently, cells were incubated with Alexa Fluor-conjugated secondary antibodies (488 nm, 555 nm, or 647 nm) for 2 hours at RT in the dark. Cell nuclei were stained with DAPI at a 1:10,000 dilution Image acquisition was performed using the Olympus DP80 microscope. 10 representative images were taken per slide. Automated quantitative analysis of stained cell cultures was performed using MetaMorph Software V7.6 (Molecular Devices, USA) via the Multi-Wavelength Cell Scoring application, where marker-positive cells were identified based on signal intensity exceeding a predefined threshold. The list of antibodies used can be found in Supplementary Methods.

## Acknowledgements

We wish to thank Carlo Sala Frigerio, Oliver Polzer, Elena Moreno-Jiménez, Inge Huitinga, Dick Swaab, and the VIB Nucleomics Core of Leuven, Belgium, for providing feedback and resources. We are grateful to the Netherlands Brain Bank, the brain donors, and their families for their invaluable contributions to this research. This work was supported by funding from Alzheimer Nederland (WE.03-2020-04) and the Friends Foundation of the Netherlands Institute of Neuroscience (Stichting Vrienden van het Herseninstituut), which was made possible by Stichting Aan Boord and Stichting Burgland. The Dutch national e-infrastructure with the support of the Dutch Research Council (Nederlandse Organisatie voor Wetenschappelijk Onderzoek, NWO) and the SURF Cooperative (EINF-1405 and EINF-4880) was used. C.P.F. receives funding from Alzheimer Nederland (WE.03-2020-07), and P.J.L. from Alzheimer Nederland (WE.03-2018-01), ZonMW Memorabel (MODEM consortium), and the Center for Urban Mental Health. P.J.L., C.P.F. and E.S. are further supported by the Gravitation project ‘Institute for Chemical Neuroscience (iCNS)’ (024.006.009), funded by the Dutch Research Council (NWO).

## Author contribution

Conceptualization, G.T., D.A., E.S.; Project supervision, E.S., C.P.F., P.J.L.; Cohort selection and stratification, G.T., A.P., S.S., L.E.D.V, J.V.; Dentate gyrus isolation, G.T., A.P., S.S.; Nuclei isolation and library preparation, G.T., A.P., S.S., S.M.P; Computational analysis, G.T., D.A.; Data analysis supervision, E.S., K.D., J.B., W.M., O.B.; Provision of fetal samples, J.A., E.A.; Provision of cell line samples, S.T., L.R.; *In situ* validation, G.T., S.S.; *In vitro* validation, S.S., E.S.M., O.R.O.; H.L.; Writing, G.T. and E.S., with input from all authors.

## Data availability statement

SnRNA-seq data will be deposited into the Gene Expression Omnibus database, with accession number to be determined upon manuscript acceptance.

Supplementary Information is available for this paper.

