## Supplementary Material for "Unique transcriptional profiles of adult human immature neurons in healthy aging, Alzheimer’s disease, and cognitive resilience"

**Supplementary Figures**

**Supplementary Figure 1**

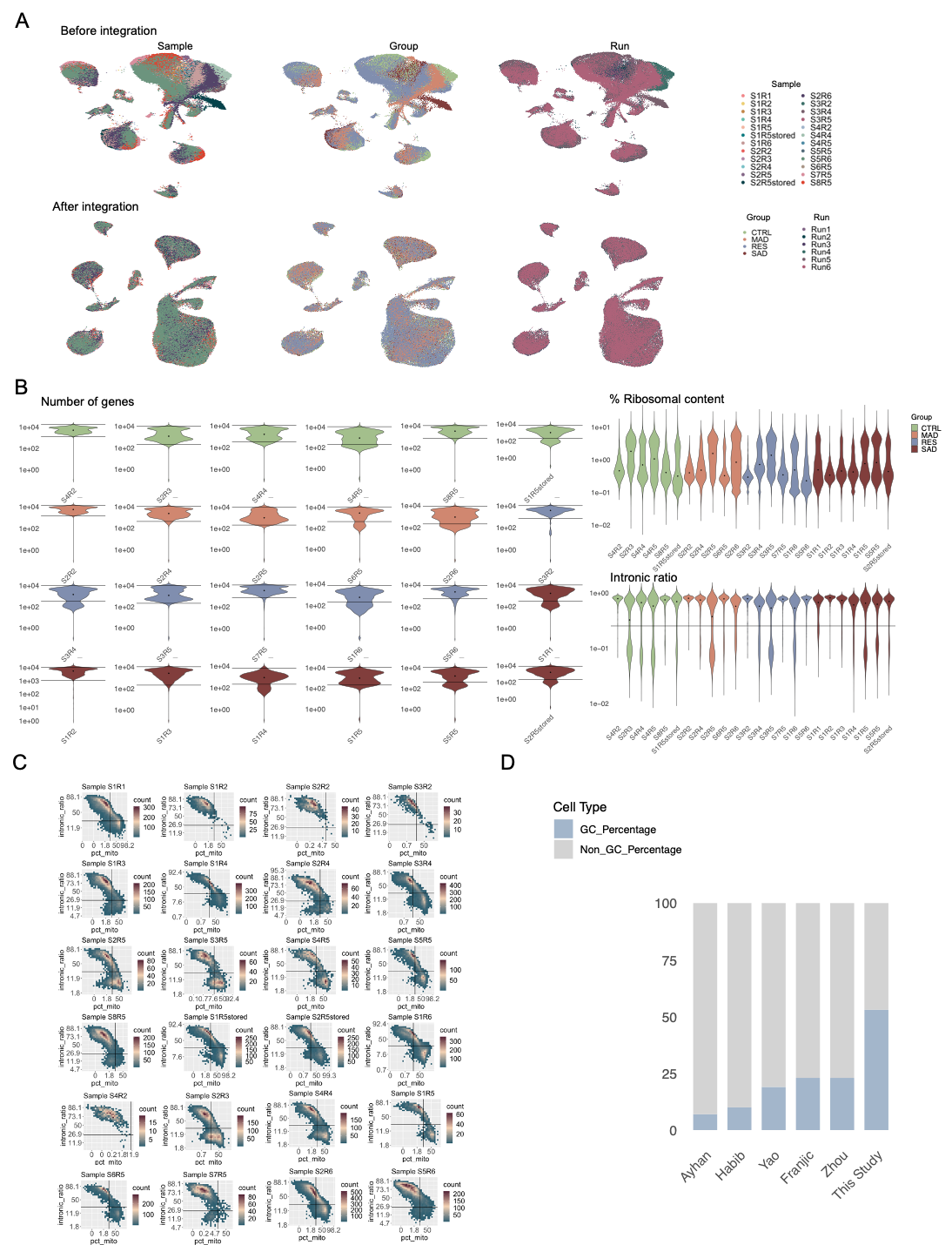

**Supplementary Figure 1| Integration and quality control metrics of human adult hippocampal niche snRNA-seq dataset. A,** Top row: UMAP plots showing data distribution before integration, colored by sample (left), group (center), and sequencing run (right). Bottom row: UMAP plots showing data distribution after integration (Harmony; variables, ‘sample’ and ‘run’), colored as in top row**. B,** Violin plots summarizing quality control metrics across samples (S), groups (color), and sequencing run (R). Left panel: Distribution of the number of genes detected per cell. Right panel: Percentage of ribosomal content (%) and intronic ratio detected per cell. Horizontal lines indicate the filtering thresholds applied. **C,** Scatter plots of intronic ratio versus mitochondrial content for individual samples. Horizontal and vertical lines represent the filtering threshold used for intronic ratio and mitochondrial content, respectively. **D,** Bar plot showing the percentage of annotated GC on the total number of cells/nuclei across previously published studies on human hippocampus.

**Supplementary Figure 2**

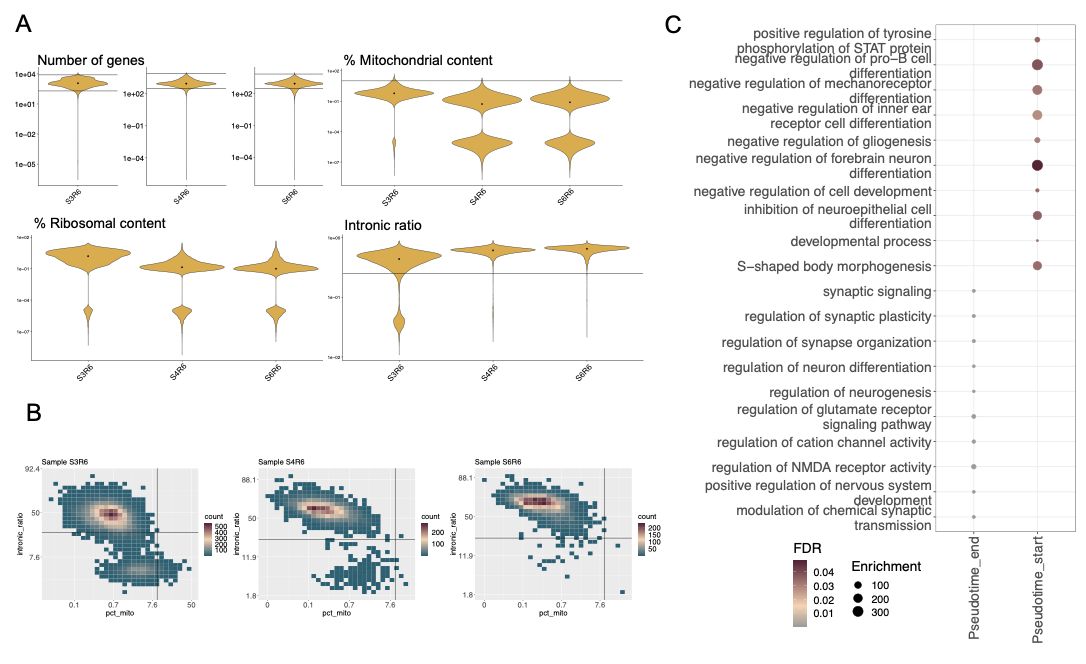
**Supplementary Figure 2| Quality control and pseudotime pathway analysis of fetal snRNA-seq. A,** Violin plots showing quality control metrics for samples. Top row: Distribution of the number of genes detected per cell and mitochondrial content (%) across samples. Bottom row: Ribosomal content (%) and intronic ratio distribution across samples. Horizontal lines indicate the filtering thresholds applied. **B,** Scatter plots of intronic ratio versus mitochondrial content for individual samples. Horizontal and vertical lines represent the filtering threshold used for intronic ratio and mitochondrial content, respectively. **C,** Dot plot of enriched gene ontology (GO) terms across pseudotime stages. GO terms enriched at the beginning and end of pseudotime trajectories are shown.

**Supplementary Figure 3**

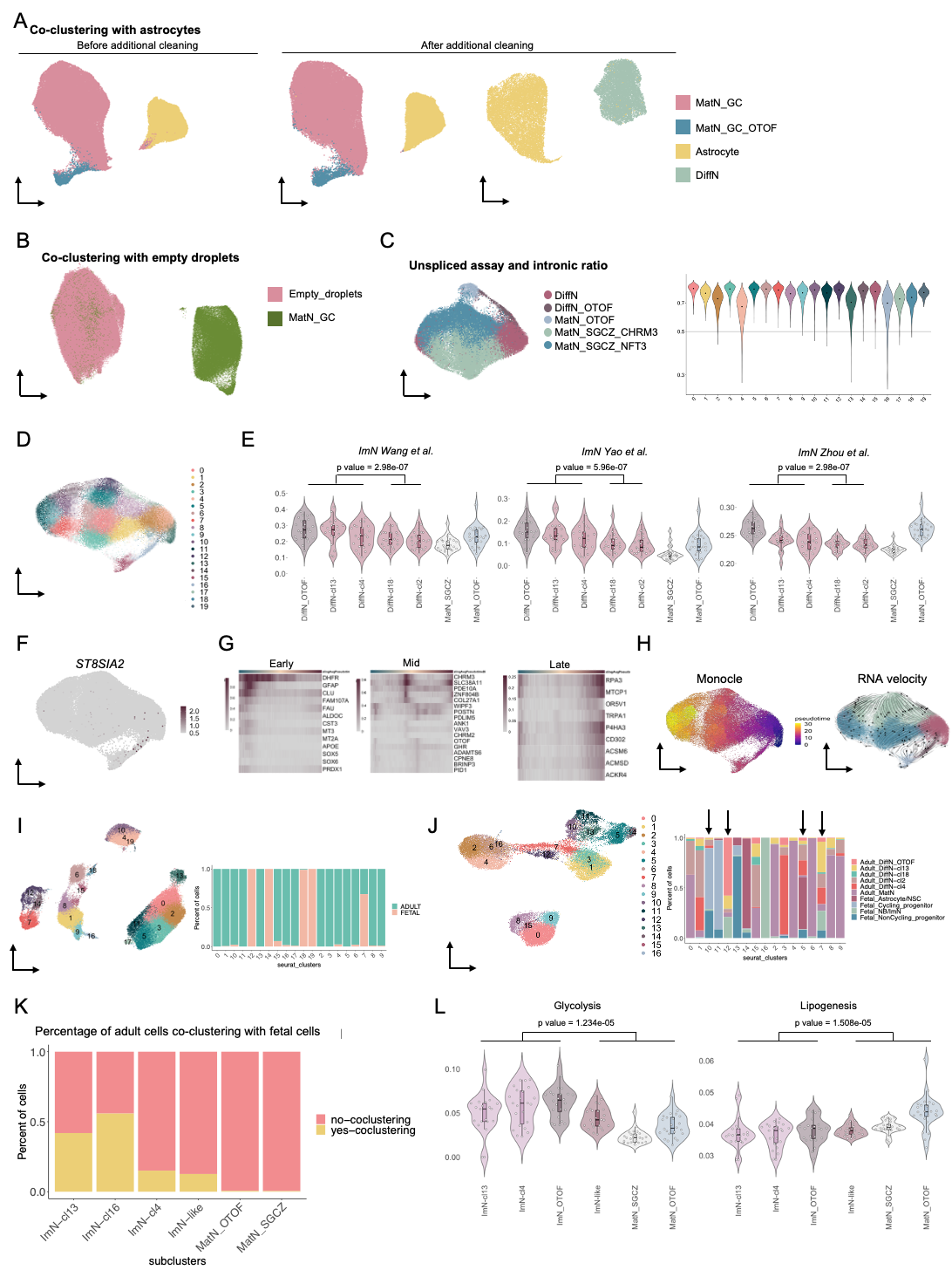

**Supplementary Figure 3| Additional quality control filtering and profiling of ImN subpopulations. A,** Subsetting and integration of GC and astrocyte clusters before and after additional quality control filtering (see panels B and C). **B,** Subsetting of the GC cluster and integration with empty/ambient RNA-containing droplets. Putative contaminated GC nuclei co-clustering with empty droplets (n = 1,386) were excluded from further analysis. **C,** UMAP plot of the GC dataset based on unspliced RNA counts, with nuclei colored by GC subpopulations as shown in Figure 3B (left). Violin plots comparing intronic ratios across GC Seurat clusters (see D), confirming robust clustering independent of cytoplasmic RNA contamination after additional QC filtering (right). **D,** UMAP visualization of the GC dataset after additional quality control filtering, displaying Seurat clusters identified at the selected resolution (=1.7)**. E,** Violin plots representing combined expression levels of gene groups defining the ImN transcriptional profile, as characterized in previous studies [ref]. Each dot represents the pseudobulk UCell score of individual donor samples (n = 24). Statistical significance between ImN (DiffN cl. 13 + DiffN cl. 4 + DiffN_OTOF) and ImN-like (DiffN cl. 2 + DiffN cl. 18) subpopulations was assessed using a one-sided Wilcoxon rank-sum test, with p-values indicated. **F,** UMAP of GC dataset showing the expression of *ST8SIA2*. **G,** Heatmap of differentially expressed genes along pseudotime inferred using Slingshot trajectory analysis. **H**, Trajectory inference analysis using Monocle and RNA velocity. **I,** UMAP plot showing the integration of adult GC and fetal neurogenic datasets to assess co-clustering (left). Barplot showing the proportions of adult and fetal nuclei by cluster. Clusters showing no co-clustering between fetal and adult in the integrated UMAP were excluded from further analysis (See Figure 3H and Supplementary Figure 3J). **J,**	UMAP plot after removing non-co-clustering adult nuclei (integrated dataset, clusters 11, 5, 3, 0, 2, 13, 17 in panel I) (left). Barplot showing the proportions of adult and fetal subpopulations by cluster (right). **K,** Percentage of adult GC nuclei co-clustering with fetal neurogenic cells, confirming immature neuronal identity within specific GC subclusters. **L,** Violin plots depicting glycolysis and lipogenesis signatures across GC subtypes. Single dots represent pseudobulk UCell score of individual donor samples (n=24). Statistical significance between ImN (DiffN cl. 13 + DiffN cl. 4 + DiffN_OTOF) and ImN-like (DiffN cl. 2 + DiffN cl. 18)+ Mature GC (MatN_SGCZ + MatN_OTOF) subpopulations was assessed using a one-sided Wilcoxon rank-sum test, with p-values indicated.

**Supplementary Figure 4**

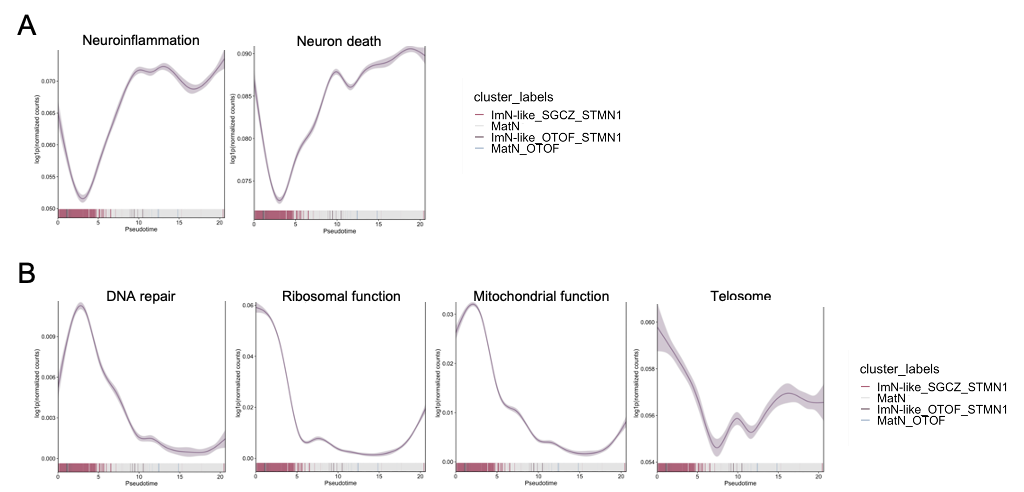

**Supplementary Figure 4| Transcriptional features of immature neurons across pseudotime. A, B,** Fitted curves showing transcriptional scores along pseudotime for dedifferentiation signatures **(A)** and cellular age-associated pathways (**B**).

**Supplementary Figure 5**

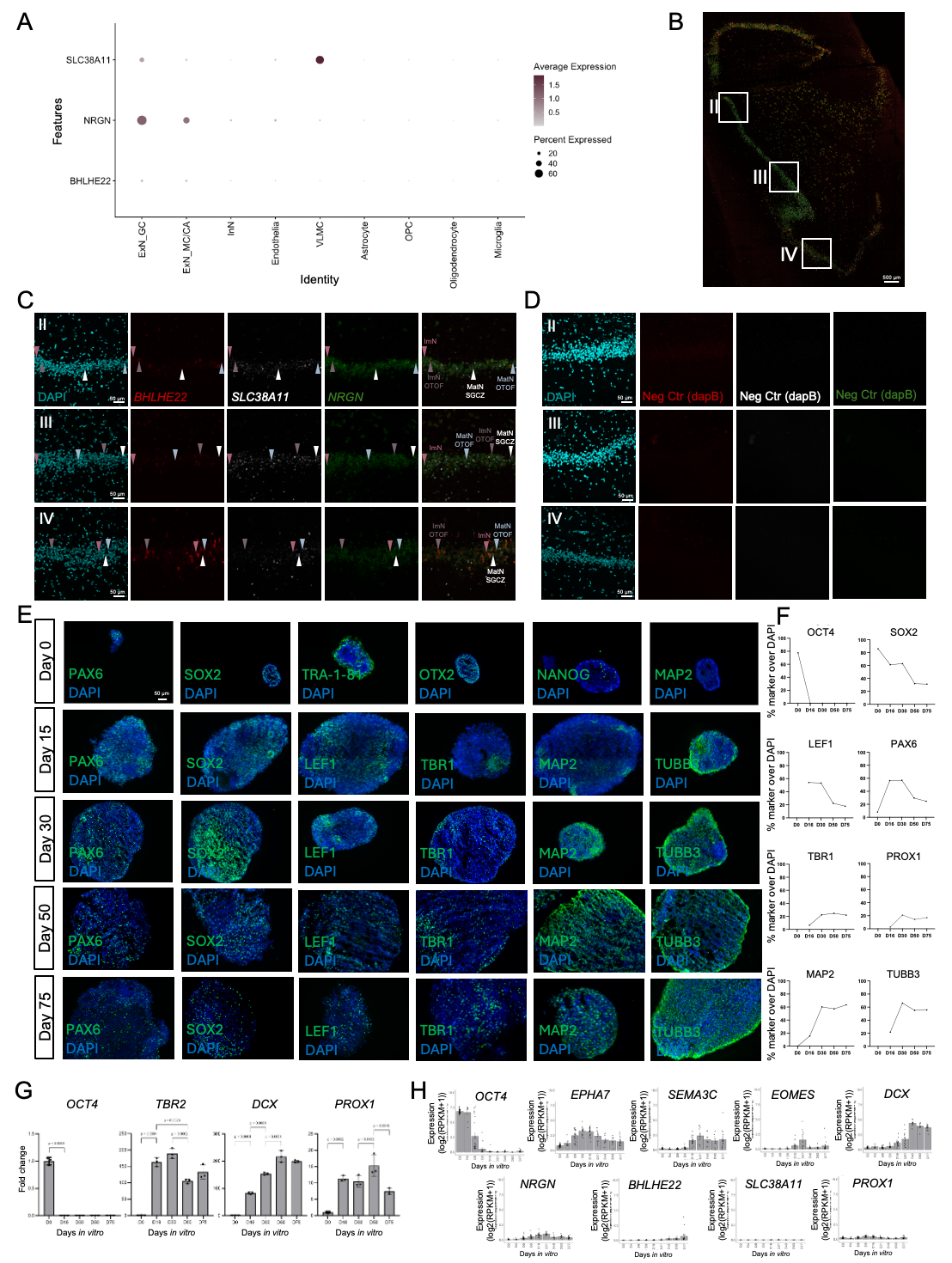

**Supplementary Figure 5|** **Validation of immature neuronal profiles in adult human hippocampus and iPSC-derived hippocampal spheroids. A,** Dot plot depicting expression profiles of selected marker genes (*NRGN*, *BHLHE22*, and *SLC38A11*) across all cell types in adult DG. **B,** Representative tile scan image of the DG showing the selected regions of interest (I-III) for subsequent analysis. White boxes indicate areas where cellular populations were quantified for marker expression**. C,** RNAScope validation of GC subpopulations in DG subregion (I-III) of an independent healthy adult donor. Arrowheads indicate one exemplary cell for each subpopulation. For color code, see Figure 5C. **D,** Negative controls for RNAScope, showing the absence of signal for markers in DG subregions I-III using a dapB probe. **E,** Immunofluorescence depicting temporal expression of developmental markers during the differentiation of human iPSC-derived hippocampal spheroids: stem cells (SOX2, PAX6, TRA-1-81, OTX2, NANOG), intermediate neural progenitors (LEF1, TBR1), neurons (MAP2, TUBB3), and GC neurons (PROX1) at various differentiation stages (Day 0 to Day 75). **F,** Quantification of marker expression in iPSC-derived hippocampal spheroids immunofluorescence images across differentiation time points. Percentage of DAPI-positive cells expressing OCT4, SOX2, LEF1, PAX6, TBR1, PROX1, MAP2, and TUBB3 is shown. **G,** Semi-quantitative real-time PCR analysis of gene expression in hippocampal spheroids during neuronal differentiation. Data are shown as mean ± standard deviation (SD). Statistical comparisons were performed using one-way ANOVA followed by Tukey's test; p-value is indicated. **H,** Marker expression in human iPSC differentiation towards cortical neurons (re-analysis PMID: 31974374).

**Supplementary Figure 6**

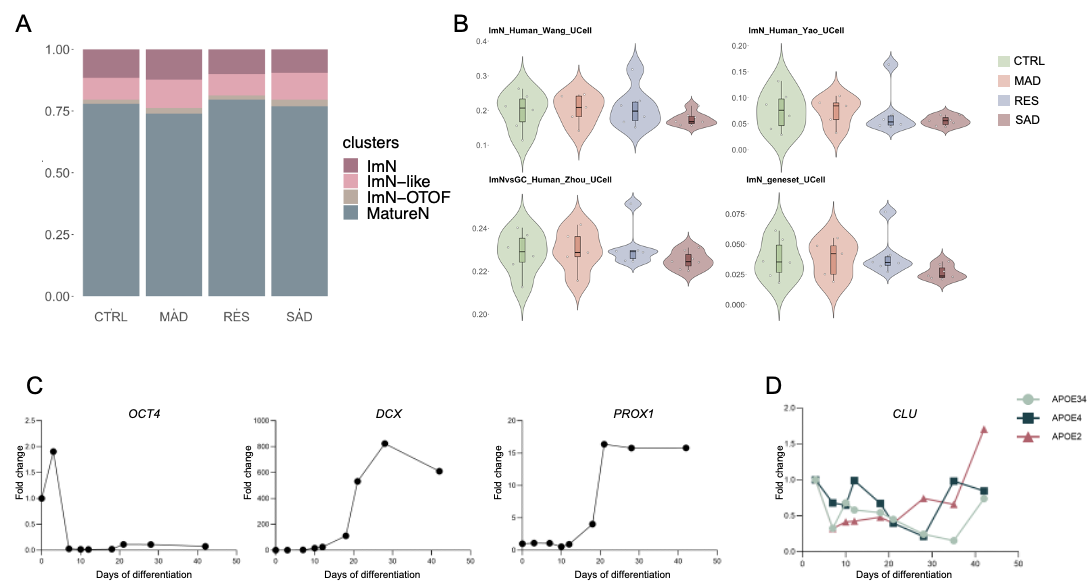

**Supplementary Figure 6| ImN gene programs across donor groups. A,** Normalized proportion of GC subpopulations across groups. **B,** Violin plots showing UCell enrichment scores for various ImN transcriptional signatures in ImN of CTRL, MAD, RES, and SAD groups. ImN signatures (Wang, Yao, Zhou) and the ImN-specific gene set from this study are depicted. Each dot corresponds to the pseudobulk UCell score of the individual donor samples in each group. **C,** Semi-quantitative real-time PCR analysis of marker gene expression (stem cells, *OCT4*; neuronal differentiation, *DCX*; GC neuron differentiation, *PROX1*) in hippocampal neurons derived from human iPSCs carrying the APOE3/4 genotype along differentiation. **D,** Semi-quantitative real-time PCR analysis of *CLU* expression in hippocampal neurons derived from human iPSCs carrying the APOE2, APOE3/4, and APOE4 genotypes during differentiation.

**Supplementary Methods**

**Primer sequences used for semi-quantitative real-time PCR.**

|  | **Forwards sequence** | **Reverse sequence** |
| --- | --- | --- |
| *GAPDH* | TCAAGAAGGTGGTGAAGCAGG | ACCAGGAAATGAGCTTGACAAA |
| *PSMB4* | TGTCACCGAAAAAGGTGTTG | AAGAGTCTATCTTTGAACTAGCCAAG |
| *BHLHE22* | GTCCCCGGGCTGCTAGTA | CGCTTTCGCCGTACTTGAG |
| *NRGN* | ACTGCTGCACCGAGAACG | GAAAACTCGCCTGGATTTTG |
| *EPHA7* | ACCCCGATACGAACATACCA | TGCAAGTTCCCAGTACTCCA |
| *SEMA3C* | GGAGGAGCCGAAGACAAGAT | GATAGATGCCTGCGGAGACT |
| *PROX1* | GAGTGCGGCGATCTTCAAG | CGTGCGTACTTCTCCATCTG |
| *SLC38A11* | TCCCGCCTTATCCATATGTCC | TGTCACAAAGCATTCCATAGGG |
| *DCX* | TGCCTCAGGGAGTGCGTTA | GAACAGACATAGCTTTCCCCTTC |
| *OCT4* | TCGAGAACCGAGTGAGAGG | GAACCACACTCGGACCACA |
| *TBR2* | TCAAATTCCACCGCCACCAA | GCAGTGGGATTGAGTCCGTT |
| *CLU* | ACAACGAGCTGCTAAAGTCC | AACGTCCGAGTCAGAAGTGT |

**List of primary antibodies used for immunostaining.**

| **Antibody** | **Species** | **Dilution** | **Manufacturer** | **Reference** |
| --- | --- | --- | --- | --- |
| OTX2 | Goat | 1:100 | R&D Systems | BAF1979 |
| NANOG | Mouse | 1:200 | ThermoFisher Scientific | MA1-017 |
| TRA-1-81 | Mouse | 1:400 | Invitrogen | 41-1100 |
| SOX2 | Goat | 1:100 | R&D Systems | AF2018 |
| TUBB3 | Mouse | 1:400 | Sigma Aldrich | T8660 |
| PAX6 | Rabbit | 1:400 | Cell Signaling | 60433S |
| LEF1 | Rabbit | 1:200 | Cell Signaling | 2230S |
| PROX1 | Rabbit | 1:100 | ThermoFisher Scientific | PA5-85552 |
| MAP2 | Chicken | 1:1000 | Abcam | Ab92434 |
| TBR1 | Rabbit | 1:400 | Sigma | AB10554 |
| OCT4 | Mouse | 1:200 | Millipore Sigma | MABD76 |

**Supplementary Tables**

**Supplementary Table 1: Overview of the human adult hippocampal dataset.** Summary of donor information, sequencing characteristics, cell type markers, and cell type abundance.

**1.1 donors_ clinfo**: Detailed clinical and pathological information of the human specimens used for the human adult hippocampal dataset

**1.2 donors_seqinfo**: Report of sequencing characteristics per specimen

**1.3 celltype_markers**: Differentially expressed genes among major cell types of human adult hippocampus

**1.4 celltype_numbers**: Numbers of the identified cell types per specimen

**Supplementary Table 2: Overview of the human fetal hippocampal dataset.** Summary of sequencing characteristics, cell type markers, neurogenic lineage gene drivers, GO terms, and cell cycle genes

**2.1 donors_seqinfo**: Report of sequencing characteristics per specimen

**2.2 celltype_markers**: Differentially expressed genes among major cell types of human fetal hippocampus

**2.3 neurog_markers**: Differentially expressed genes among cell types of human fetal neurogenic lineage

**2.4 DEG_trajectory**: Differentially expressed genes along the inferred neurogenic lineage trajectory

**2.5 GO_trajectory**: List of GO terms associated with genes differentially expressed along the neurogenic trajectory. Highlighted terms are represented in Figure 2E

**2.6 genes_cellcycle**: List of genes used for inferring cell cycle, as reported by *Schwabe et al.*

**Supplementary Table 3: Overview of adult GC subset data.** GC subtype markers, ambient RNA markers, gene sets used for UCell scoring, and the proportions of ImN across studies.

**3.1 GC_markers:** Differentially expressed genes across subpopulations of the human adult GC subset. For DiffN subpopulations, markers are calculated for pooled subclusters (DiffN) as well as for individual subclusters (2, 4, 13, 18).

**3.2 ambientRNA_markers:** Differentially expressed genes between annotated GC droplets and empty droplets, representing markers for ambient RNA contamination.

**3.3 genes_Ucell:** Gene sets applied for UCell scoring to assess cellular states in Figure 3D and Supplementary Figure 3E,L.

**3.4 genes_ImN:** Curated list of widely used ImN markers for UCell scoring in Figure 3G, along with their average expression across GC subpopulations.

**3.5 proportions_ImN:** Proportions of ImN across previously published studies.

**Supplementary Table 4: ImN transcriptional identity.** Gene sets, differential expression, and GO analysis related to ImN age and identity.

**4.1 genes_Ucell:** Gene sets applied for UCell scoring to assess different transcriptional signatures in Figure 4B,C.

**4.2 DEG_AdultFetal**: Differentially expressed genes between adult (control only) ImN and fetal ImN/NB.

**4.3 GO_AdultFetal:** List of GO terms associated with genes differentially expressed between adult control ImN and fetal ImN/NB.

**4.4 DEG_ImNMatN**: Differentially expressed genes between adult ImN (ImN+ImN_OTOF) and adult MatN (MatN_SGCZ + MatN_OTOF)

**4.5 GO_ ImNMatN:** List of GO terms associated with genes differentially expressed between adult ImN (ImN+ImN_OTOF) and adult MatN (MatN_SGCZ + MatN_OTOF). Highlighted terms are represented in Figure 4E.

**Supplementary Table 5: Validation of ImN *in situ.*** DEG used for RNAScope probe selection and quantification analysis.

**5.1 DEG_ ImNvsMatN+Astro**: Differentially expressed genes between ImN and MatN+Astro to identify ImN-specific markers.

**5.2 DEG_ MatNvsImN+ImN-like**: Differentially expressed genes between MatN and ImN+ImN-like to identify MatN-specific markers.

**5.3 QuantRNAScope**: Quantification analysis of selected transcript per nucleus in postmortem control human hippocampus. Highlighted values (normalized expression across images I,II,III, IV) are showed in Figure 5D.

**Supplementary Table 6: DEG and cell-cell interaction differences across groups.** DEG of ImN across groups, DEG of other GC subtypes across groups, CellPhoneDB analysis.

**6.1 DEG_ImNgroups:** Differentially expressed genes of ImN among groups. DEG are calculated for pooled ImN (cl4+cl13+ImN_OTOF) as well as for individual ImN subclusters (ImN cl 4, ImN cl 13, ImN_OTOF ). Genes highlighted are reported in Figure 6C.

**6.2 DEG_GCgroups:** Differentially expressed genes of ImN-like and MatN GC subpopulations among groups.

**6.3 Number_CellPhone:** Number of unique intercellular interactions as visualized in Figure 6D.

**6.4 All_CellPhone:** All intercellular interactions as calculated by CellphoneDB, unique interactions used for Figure 6D are indicated. Interactions visualized in Figure 6E are highlighted.
